# The large GTPase Sey1/atlastin mediates lipid droplet- and FadL-dependent intracellular fatty acid metabolism of *Legionella pneumophila*

**DOI:** 10.1101/2022.12.05.519141

**Authors:** Dario Hüsler, Pia Stauffer, Bernhard Keller, Desirée Böck, Thomas Steiner, Anne Ostrzinski, Bianca Striednig, A. Leoni Swart, François Letourneur, Sandra Maaß, Dörte Becher, Wolfgang Eisenreich, Martin Pilhofer, Hubert Hilbi

**Affiliations:** Institute of Medical Microbiology, University of Zurich, Gloriastrasse 30/32, 8006 Zurich, Switzerland; Institute of Molecular Biology & Biophysics, ETH Zurich, Otto-Stern-Weg 5, 8093 Zurich, Switzerland; Bavarian NMR Center – Structural Membrane Biochemistry, School of Natural Sciences, Technical University of Munich, Lichtenbergstrassse 4, 85747 Garching, Germany; Institute of Microbiology, University of Greifswald, Felix-Hausdorff-Strasse 8, 17489 Greifswald, Germany; UMR5235, DIMNP, CNRS/Université Montpellier, Place Eugène Bataillon, Montpellier, 34095, Cedex 5, France

**Keywords:** Amoeba, atlastin, *Dictyostelium discoideum*, fatty acid transport, large fusion GTPase, lipid droplet, host-pathogen interaction, *Legionella pneumophila*, Legionnaires’ disease, pathogen vacuole.

## Abstract

The facultative intracellular bacterium *Legionella pneumophila* employs the Icm/Dot type IV secretion system (T4SS) to replicate in a unique membrane-bound compartment, the *Legionella*-containing vacuole (LCV). The endoplasmic reticulum (ER)-resident large fusion GTPase Sey1/atlastin promotes remodeling and expansion of LCVs, and the GTPase is also implicated in the formation of ER-derived lipid droplets (LDs). Here we show that LCVs intimately interact with palmitate-induced LDs in *Dictyostelium discoideum* amoeba. Comparative proteomics of LDs isolated from the *D. discoideum* parental strain Ax3 or Δ*sey1* revealed 144 differentially produced proteins, of which 7 or 22 were exclusively detected in LDs isolated from strain Ax3 or Δ*sey1*, respectively. Using dually fluorescence-labeled amoeba producing the LCV marker P4C-GFP or AmtA-GFP and the LD marker mCherry-perilipin, we discovered that Sey1 and the *L. pneumophila* Icm/Dot T4SS as well as the effector LegG1 promote LCV-LD interactions. *In vitro* reconstitution of the LCV-LD interactions using purified LCVs and LDs from *D. discoideum* Ax3 or Δ*sey1* revealed that Sey1 and GTP promote this process. The LCV-LD interactions were impaired for Δ*sey1*-derived LDs, suggesting that Sey1 regulates LD composition. Palmitate promoted the growth of (i) *L. pneumophila* wild-type in *D. discoideum* Ax3 but not in Δ*sey1* mutant amoeba and (ii) *L. pneumophila* wild-type but not Δ*fadL* mutant bacteria lacking a homologue of the *E. coli* fatty acid transporter FadL. Finally, isotopologue profiling indicated that intracellular *L. pneumophila* metabolizes ^13^C-palmitate, and its catabolism was reduced in *D. discoideum* Δ*sey1* and *L. pneumophila* Δ*fadL*. Taken together, our results reveal that Sey1 mediates LD- and FadL-dependent fatty acid metabolism of intracellular *L. pneumophila*.

## Introduction

The causative agent of Legionnaires’ disease, *Legionella pneumophila*, is a facultative intracellular bacterium, which adopts a similar mechanism to replicate in free-living protozoa and lung macrophages (Newton et al., 2010; Boamah et al., 2017; Mondino et al., 2020). To determine the interactions with eukaryotic host cells, *Legionella* spp. employ the genus-conserved Icm/Dot type IV secretion system (T4SS), which in *L. pneumophila* translocates more than 330 different “effector” proteins (Qiu and Luo, 2017; Hilbi and Buchrieser, 2022; Lockwood et al., 2022). The effector proteins subvert pivotal processes and establish a unique replication niche, the *Legionella*-containing vacuole (LCV), which communicates with the endosomal, secretory and retrograde vesicle trafficking pathway, but restricts fusion with lysosomes (Isberg et al., 2009; Asrat et al., 2014; Finsel and Hilbi, 2015; Personnic et al., 2016; Sherwood and Roy, 2016; Bärlocher et al., 2017; Steiner et al., 2018a; Swart and Hilbi, 2020). Given that many cellular pathways and effector protein targets are conserved, the genetically tractable amoeba *Dictyostelium discoideum* is a versatile and powerful model to analyze pathogen-phagocyte interactions (Cardenal-Munoz et al., 2017; Swart et al., 2018).

Peculiarities of LCV formation are the phosphoinositide (PI) lipid conversion from PtdIns(3)*P* to PtdIns(4)*P* (Weber et al., 2006; Weber et al., 2014; Steiner et al., 2018a; Weber et al., 2018; Swart and Hilbi, 2020), interception of and fusion with endoplasmic reticulum (ER)-derived vesicles (Kagan and Roy, 2002; Robinson and Roy, 2006; Arasaki et al., 2012) and a tight association with the ER (Swanson and Isberg, 1995; Robinson and Roy, 2006), which persists even upon isolation and purification of intact LCVs (Urwyler et al., 2009; Hoffmann et al., 2014; Schmölders et al., 2017). Indeed, LCVs form extended membrane contact sites (MCSs) with the ER (https://www.biorxiv.org/content/10.1101/2022.06.17.496549v1). Using dually fluorescence-labeled *D. discoideum* and defined deletion mutant strains, we recently revealed that the PtdIns(4)*P* 4-phosphatase Sac1 and VAMP-associated protein (Vap) localize to the ER, but not to the LCV membrane, while the putative lipid transport proteins oxysterol binding protein 8 (OSBP8) and OSBP11 specifically localized to the ER or the LCV membrane, respectively. Moreover, these MCS components were implicated in LCV remodeling and intracellular growth of *L. pneumophila*.

The ER is a highly dynamic organelle (Shibata et al., 2006; Hu and Rapoport, 2016; Nixon-Abell et al., 2016), and its morphology and dynamics are largely controlled by the reticulon family of membrane tubule-forming proteins (Voeltz et al., 2006; Hu et al., 2008) and the atlastin family of trans-membrane large fusion GTPases (Hu et al., 2009; Orso et al., 2009). Atlastins are conserved from yeast to plants and mammals (Anwar et al., 2012; Zhang et al., 2013), mediate the homotypic fusion of ER tubules and share a similar domain organization, which comprises an N-terminal GTPase domain linked through a helical bundle (HB) domain to two adjacent transmembrane segments and a C-terminal tail that contains an amphipathic helix (Hu and Rapoport, 2016). Structural and biochemical studies revealed that upon GTP binding, the GTPase and HB domains of atlastins on two distinct ER tubules dimerize, and the trans-homodimers pull together opposing membranes thus facilitating their fusion (Bian et al., 2011; Byrnes and Sondermann, 2011; Liu et al., 2012; Byrnes et al., 2013; Liu et al., 2015).

*D. discoideum* produces a single orthologue of human atlastin 1-3 (Atl1-3) termed Sey1, which shares the same domain organization as the mammalian atlastins (Steiner et al., 2017). Initially, Sey1, Atl3 and reticulon-4 (Rtn4) were identified by proteomics in intact LCVs purified from *L. pneumophila*-infected *D. discoideum* or macrophages, respectively (Hoffmann et al., 2014), and the localization of Sey1/Atl3 and Rtn4 to LCVs was validated by fluorescence microscopy (Steiner et al., 2017). Sey1 is not implicated in the formation of the PtdIns(4)*P*-positive LCV membrane and not essential for the recruitment of ER, but promotes pathogen vacuole expansion and enhances intracellular replication of *L. pneumophila* (Steiner et al., 2017; Steiner et al., 2018b). The production of a catalytically inactive, dominant-negative Sey1_K154A mutant protein, or the depletion of mammalian Atl3, restricts *L. pneumophila* replication and impairs LCV maturation. *D. discoideum* Δ*sey1* mutant amoeba are enlarged but grow and develop similarly to the parental strain (Hüsler et al., 2021). The mutant strain shows pleiotropic defects, including aberrant ER architecture and dynamics, inability to cope with prolonged ER stress, defective intracellular proteolysis, cell motility and growth on bacterial lawns (Hüsler et al., 2021). Moreover, in the Δ*sey1* mutant strain LCV-ER interactions, LCV expansion and intracellular *L. pneumophila* replication is impaired. Taken together, Sey1/Atl3 controls circumferential ER remodeling during LCV maturation and intracellular replication of *L. pneumophila* (Steiner et al., 2017; Steiner et al., 2018b; Hüsler et al., 2021).

In addition to promoting ER dynamics, atlastins contribute to a number of other cellular processes including the biogenesis of ER-derived lipid droplets (LDs) (Klemm et al., 2013). LDs are the major cellular storage compartments of neutral lipids; however, they are also involved in many other cellular processes such as energy homeostasis, lipid metabolism, generation of membrane lipids and signaling molecules as well as retention of harmful proteins and lipids (Walther and Farese, 2012; Hashemi and Goodman, 2015; Welte, 2015; Kimmel and Sztalryd, 2016; Welte and Gould, 2017). In *D. discoideum*, LDs accumulate upon feeding the cells with fatty acids (in particular palmitate) or bacteria (Du et al., 2013). Proteomics and lipidomics analysis of *D. discoideum* LDs revealed that the lipid constituents are similar to mammalian LDs and comprise mainly triacylglycerol (57%), free fatty acids (22 %) and sterol esters (4%). LDs are coated by a polar phospholipid monolayer and distinct proteins (Du et al., 2013), such as perilipin (Miura et al., 2002), and small GTPases, as well as ER proteins (reticulon C, RtnlC; protein disulfide isomerase, PDI; lipid droplet membrane protein, LdpA; and 15 lipid metabolism enzymes), the latter reflecting their cellular organelle origin (Du et al., 2013). LDs are transported along microtubules and actin filaments or moved by actin polymerization (Welte, 2004, 2009; Pfisterer et al., 2017; Welte and Gould, 2017; Kilwein and Welte, 2019), and they form contact sites with various cell organelles (Kumar et al., 2018; Benador et al., 2019; Yeshaw et al., 2019; Herker et al., 2021).

Only after the replication-permissive LCV has been formed, *L. pneumophila* engages in intracellular replication. The bacteria employ a biphasic lifestyle comprising a transmissive (motile, virulent) and a replicative phase (Molofsky and Swanson, 2004). *L. pneumophila* is an obligate aerobe, which previously has been thought to rely on certain amino acids as carbon and energy source (Abu Kwaik and Bumann, 2013; Manske and Hilbi, 2014). Indeed, isotopologue profiling studies with stable ^13^C-isotopes indicated that serine is a major carbon and energy source for *L. pneumophila* and readily metabolized by the bacteria (Eylert et al., 2010). More recent physiological and isotopologue profiling studies established that glucose, inositol and glycerol are metabolized by *L. pneumophila* under extracellular and intracellular conditions (Eylert et al., 2010; Harada et al., 2010; Häuslein et al., 2016; Manske et al., 2016). Finally, isotopologue profiling studies indicated that extracellular *L. pneumophila* also efficiently catabolizes exogenous [1,2,3,4-^13^C_4_]palmitic acid, yielding ^13^C_2_-acetyl-CoA, which is used to synthesize the storage compound poly-hydroxybutyrate (PHB) (Häuslein et al., 2017).

It is unknown how fatty acids are taken up by *L. pneumophila*. In *E. coli*, the long-chain fatty acid transporter FadL localizes to the outer membrane, where the monomeric protein adopts a 14-stranded, anti-parallel β barrel structure (van den Berg et al., 2004). The N-terminal 42 amino acid residues of FadL form a small ‘hatch’ domain that plugs the barrel, and the hydrophobic substrate leaves the transporter by lateral diffusion into the outer membrane (Hearn et al., 2009).

*L. pneumophila* encodes a homolog of *E. coli* FadL, Lpg1810, which was identified as a surface-associated protein by fluorescence-labeling and subsequent mass spectrometry (MS), confirming its presence in the outer membrane (Khemiri et al., 2008).

Given the role of LDs as lipid storage organelles regulated by atlastins, we set out to analyze the contribution of LDs, Sey1 and FadL for intracellular replication and palmitate catabolism of *L. pneumophila* in *D. discoideum*. We found that Sey1 regulates LD protein composition and promotes Icm/Dot-dependent LCV-LD interactions as well as FadL-dependent fatty acid metabolism of intracellular *L. pneumophila*.

## Results

### LCV interaction and protein composition of *D. discoideum* LDs

To initially explore whether fatty acids and/or LDs play a role for intracellular replication of *L. pneumophila*, we fed *D. discoideum* with palmitate, and assessed intracellular replication of the bacteria. Feeding with 200 µM palmitate overnight significantly promoted the intracellular growth of *L. pneumophila*, while higher concentrations of palmitate had a negative effect on growth (**Fig. S1A**). To assess whether the growth-promoting effect of palmitate might involve LDs, we thought to visualize the possible interactions between LCVs and LDs. To this end, *D. discoideum* strain Ax3 was fed with palmitate, infected with the *L. pneumophila* wild-type strain JR32 and subjected to cryo-electron tomography (cryoET). The obtained cryotomograms clearly show an extensive interaction between LCVs and LDs (**Fig. 1A**). Upon contact with the LCV, the LDs tightly adhere to the limiting membrane of the pathogen vacuole, followed by what appears to be an integration or even dissolution of the limiting membrane. Hence, LCVs undergo robust and intimate interactions with LDs in infected *D. discoideum*.

**Figure 1.**
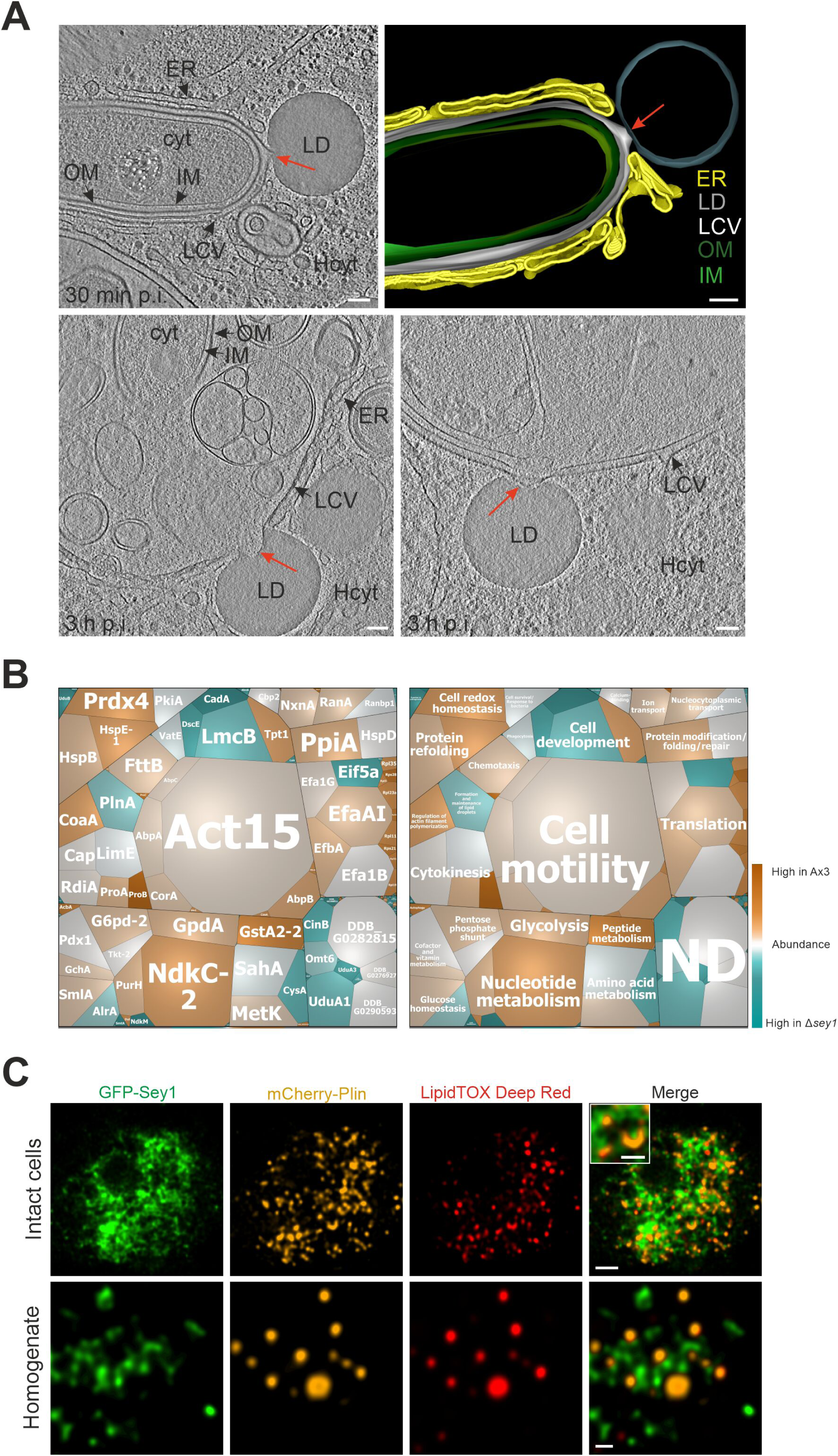
LCVs interact intimately with lipid droplets harboring Sey1, RanA and RanBP1. (**A**) Representative cryotomograms of *D. discoideum* Ax3, fed (3 h) with 200 µM sodium palmitate and infected (MOI 100) with *L. pneumophila* wild-type (JR32) for 30 min (**top**) or 3 h (**bottom**). Intimate LCV-LD interactions are clearly visible (**red arrows**). Reconstruction of LCV-LD interaction observed at 30 min p.i. (**top; right**). OM, outer membrane; IM, inner membrane; LCV, *Legionella*-containing vacuole (limiting membrane); ER, endoplasmic reticulum; LD, lipid droplet; cyt, *L. pneumophila* cytoplasm; Hcyt, host cell cytoplasm. Scale bars: 100 nm. (**B**) Voronoi treemaps displaying abundance of proteins on isolated lipid droplets from *D. discoideum* Ax3 and Δ*sey1*. The treemaps were generated by taking into account proteins exclusively identified on LDs from Ax3 or Δ*sey1*, or proteins showing significant difference in abundance between the two samples. Additionally, the 50 most abundant proteins in each sample were also included. The mosaic tiles are represented as (**left**) single proteins or (**right**) grouped in functional domains, which in turn are assembled according to general function (black frames). Individual iBAQ values of proteins on *D. discoideum* Ax3 LDs are represented by the area of the mosaic tiles (large tiles: more abundant, small tiles: less abundant). Comparison of abundances of proteins on Ax3 LDs with Δ*sey1* LDs is represented by the color gradient of the mosaic tiles (orange: more abundant on Ax3 LDs, grey: equally abundant on Ax3 and Δ*sey1* LDs, blue: more abundant on Δ*sey1* LDs). ND, Not Defined. Representative data for three independent experiments. (**C**) Representative fluorescence micrographs of intact (**top**) or homogenized (**bottom**) *D. discoideum* Ax3 producing GFP-Sey1 (pBS001) and mCherry-Plin (pHK102), fed overnight with 200 µM sodium palmitate and stained with LipidTOX Deep Red. Scale bars: intact cells (2 µm, inset: 1 µm), homogenate (0.5 µm).

Given that Sey1/atlastin is implicated in LD formation, we next assessed the role of Sey1 for the number, size and composition of LDs in *D. discoideum*. Dually fluorescence-labeled *D. discoideum* producing the LD marker mCherry-Plin and the phagosome marker AmtA-GFP were fed with 200 µM palmitate overnight, fixed and stained with LipidTOX Deep Red, and the number and size of LDs were quantified (**Fig. S1BC**). The feeding with palmitate indeed resulted in an 8-fold increase of LDs per cell, but neither the number nor the size of LDs were affected by the presence or absence of Sey1.

To gain further insights into the possible role of Sey1 for LD formation, we performed a comparative proteomics analysis by tandem MS of LDs isolated from *D. discoideum* Ax3 or Δ*sey1* mutant amoeba. This approach revealed 144 differentially produced proteins (log_2_ fold change > |0.8|), including some enzymes implicated in lipid metabolism (phospholipase PldA, phosphatidylinositol phosphate kinase Pik6/PIPkinA, sterol methyl transferase SmtA, acetoacetyl-CoA hydrolase) (**Fig. 1B**, **Table S1**). Among the differentially produced proteins, 7 or 22 were exclusively detected in LDs isolated from strain Ax3 or Δ*sey1*, respectively. Sey1 was identified on LDs isolated from strain Ax3, but as expected not on LDs isolated from Δ*sey1* mutant amoeba. Contrarily, the phospholipase PldA, the ER protein calnexin (CnxA) and the protein SCFD1/SLY1 implicated in ER to Golgi transport were identified only on LDs isolated from the Δ*sey1* strain (**Table S1**). The 50 most highly abundant proteins, which were not significantly different on LDs isolated from Ax3 or Δ*sey1* mutant amoeba, included perilipin (PlnA), which is involved in the formation and maintenance of LDs (Du et al., 2013), as well as the small GTPase RanA (Du et al., 2013) and its effector RanBP1 (**Table S1**). RanA is activated in *L. pneumophila*-infected cells and implicated in microtubule stabilization and LCV motility (Rothmeier et al., 2013; Swart et al., 2020b).

To assess the localization of Sey1 with regard to LDs, we used dually labeled *D. discoideum* Ax3 producing GFP-Sey1 as well as the LD marker mCherry-Plin, and further stained LDs with LipidTOX Deep Red (**Fig. 1C**). Under the conditions used, GFP-Sey1 accumulated in the vicinity of LDs in intact cells as well as in cell homogenates, but apparently did not co-localize with LDs. This staining pattern suggests that Sey1 localizes only in very low amounts to LDs, or that localization of Sey1 to LDs is impaired due to the fluorescent protein tag. To validate the localization of RanA and RanBP1 to LDs, we used dually labeled *D. discoideum* Ax3 producing either RanA-mCherry or RanBP1-GFP and GFP-Plin or mCherry-Plin, respectively, and further stained LDs with LipidTOX Deep Red (**Fig. S2A**). Under these conditions, ectopically produced RanA-mCherry or RanBP1-GFP localized to membranous structures in the cell, including to Plin- and LipidTOX Deep Red-positive LDs. In summary, comparative proteomics of LDs isolated from *D. discoideum* Ax3 or Δ*sey1* revealed that Sey1 and the phospholipase PldA were present exclusively in the parental or the mutant strain, respectively, and Plin, RanA as well as RanBP1 were detected on LDs from both *D. discoideum* strains.

### Dynamics of early LCV-LD interactions in *D. discoideum*

To assess the dynamics of early LCV-LD interactions, we used dually fluorescence-labeled *D. discoideum* producing the LCV marker P4C-GFP and the LD marker mCherry-Plin. The *D. discoideum* parental strain Ax3 or Δ*sey1* mutant amoeba were fed overnight with 200 µM palmitate, stained with LipidTOX Deep Red and infected with mCerulean-producing *L. pneumophila* JR32. Within the first hour of infection, the dynamic interactions of single LCVs with LDs were recorded for 60 s each at different time points (**Fig. 2A**, **Fig. S2B**). As the LCVs matured over the course of 1 h post infection (p.i.), the overall LCV-LD contacts gradually increased in *D. discoideum* Ax3, while they remained lower in Δ*sey1* mutant amoeba (**Fig. 2B**). Moreover, the retention time of individual LDs on LCVs was also signficantly higher in strain Ax3 than in Δ*sey1* mutant amoeba (**Fig. 2C**). Taken together, these real-time data indicate that Sey1 promotes the dynamics of LCV-LD interactions during the course of LCV maturation.

**Figure 2.**
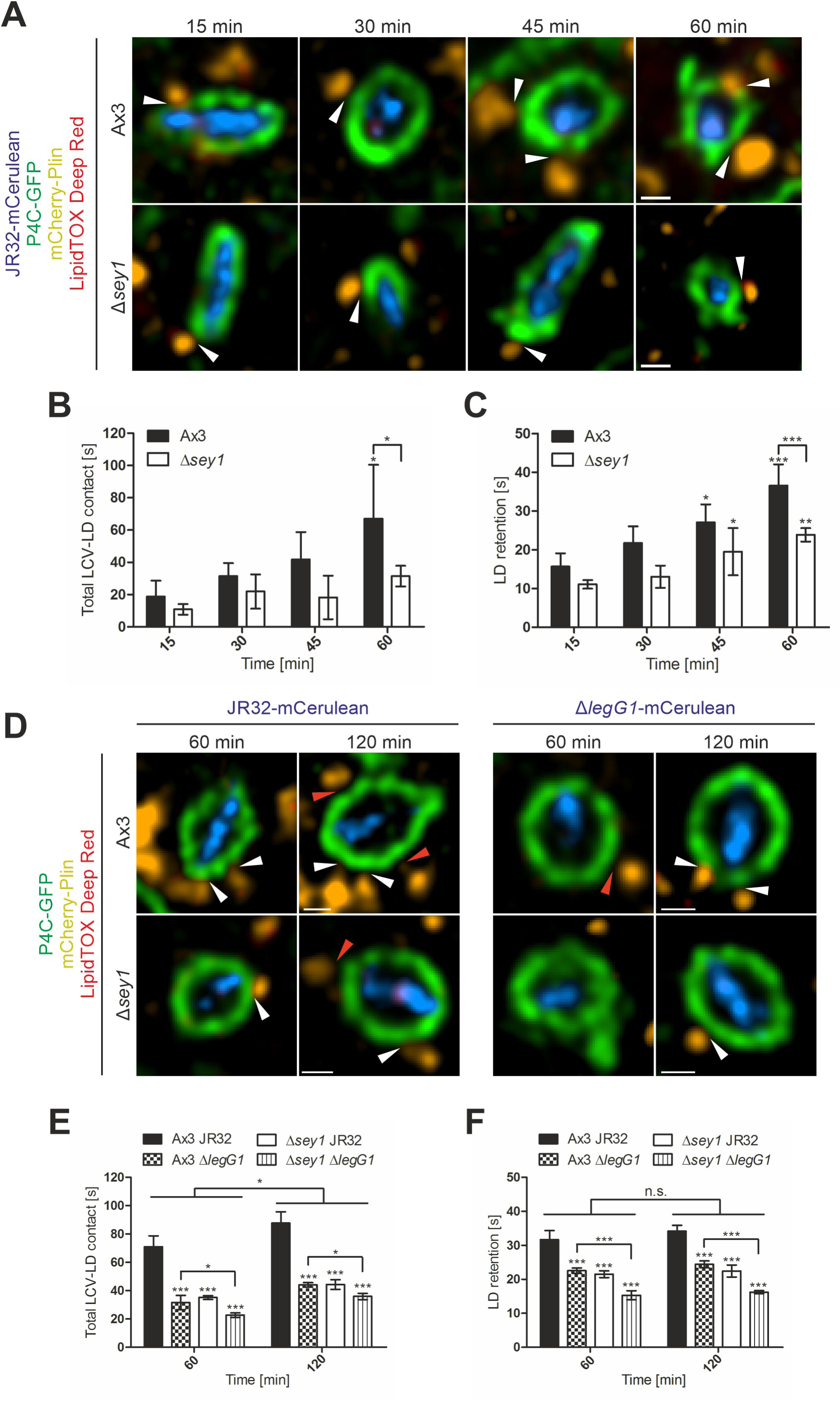
Dynamics of early LCV-LD interactions in *D. discoideum*. (**A**) Representative fluorescence micrographs of *D. discoideum* Ax3 or Δ*sey1* producing P4C-GFP (pWS034) and mCherry-Plin (pHK102), fed overnight with 200 µM sodium palmitate, stained with LipidTOX Deep Red and infected (MOI 5) with mCerulean-producing *L. pneumophila* JR32 (pNP99). Infected cells were recorded for 60 s each at the times indicated using a Leica TCS SP8. Examples are shown for contact between LDs and the LCV membrane (**white arrowheads**). Scale bars: 0.5 µm. (**B**) Quantification of (A), total contact time of LDs with the LCV recorded for 60 s at the indicated time points p.i. (video frames, n=228). Data represent means ± SD of three independent experiments (\**p* < 0.05). (**C**) Quantification of (A), retention time of single LDs with the LCV recorded for 60 s at the indicated time points p.i. (video frames, n=228). Data represent means ± SD of three independent experiments (\**p* < 0.05, \*\**p* < 0.01; \*\*\**p* < 0.001). (**D**) Representative fluorescence micrographs of *D. discoideum* Ax3 or Δ*sey1* producing P4C-GFP (pWS034) and mCherry-Plin (pHK102), fed overnight with 200 µM sodium palmitate, stained with LipidTOX Deep Red and infected (MOI 5) with mCerulean-producing *L. pneumophila* JR32 or Δ*legG1* (pNP99). Infected cells were recorded for 60 s each at the times indicated using a Leica TCS SP8. Examples are shown for contact between LDs and the LCV membrane (**white arrowheads**) or no contact (**red arrowheads**). Scale bars: 0.5 µm. (**E**) Quantification of (D), total contact time of LDs with the LCV recorded for 60 s at the indicated time points p.i. (video frames, n=228). Data represent means ± SD of three independent experiments (\**p* < 0.05, \*\*\**p* < 0.001). (**F**) Quantification of (D), retention time of single LDs with the LCV recorded for 60 s at for indicated time points p.i. (video frames, n=228). Data represent means ± SD of three independent experiments (n.s., not significant; \*\*\**p* < 0.001).

Next, we sought to identify *L. pneumophila* effector proteins, which possibly determine LCV-LD interactions. The RCC1 repeat domain effector LegG1 activates the small GTPase Ran, which in its active, GTP-bound form interacts with RanBP1 and promotes microtubule stabilization (Rothmeier et al., 2013; Swart et al., 2020b). Since we found that LDs harbor Ran and RanBP1 (**Table S1**), we tested the hypothesis that LegG1 is implicated in LCV-LD dynamics. To this end, we infected palmitate-fed *D. discoideum* Ax3 or Δ*sey1* producing P4C-GFP and mCherry-Plin with mCerulean-producing *L. pneumophila* JR32 or Δ*legG1* and additionally stained LDs with LipidTOX Deep Red (**Fig. 2D**, **Supplementary Movies S1-4**). At 1 h or 2 h p.i. the overall LCV-LD contacts were lowered by ca. 50% upon infection with Δ*legG1* (compared to JR32) or in Δ*sey1* mutant *D. discoideum* (compared to strain Ax3) (**Fig. 2E**). Intriguingly, the overall LCV-LD contacts were further significantly reduced upon infection of Δ*sey1* mutant amoeba with Δ*legG1* mutant bacteria (**Fig. 2E**). Similar results were obtained by quantifying the retention time of individual LDs on LCVs (**Fig. 2F**). In summary, these results indicate that the host large GTPase Sey1, as well as the *L. pneumophila* effector protein LegG1 promote and additively affect the dynamics of LCV-LD interactions.

### Sey1 promotes LD recruitment to LCVs in *D. discoideum*

To validate that Sey1 promotes LCV-LD interactions in palmitate-fed, fixed *D. discoideum* and to test whether the process is dependent on the *L. pneumophila* Icm/Dot T4SS, we used dually fluorescence-labeled amoeba producing mCherry-Plin and AmtA-GFP, a probe localizing to vacuoles containing either wild-type *L. pneumophila* or Δ*icmT* mutant bacteria lacking a functional T4SS (**Fig. 3A**). This approach indicated that the mean number of LDs localizing to LCVs harboring wild-type *L. pneumophila* was more than twice as high in *D. discoideum* Ax3 as compared to the Δ*sey1* mutant amoeba (**Fig. 3B**), and the effect was of similar magnitude, when the number of LDs per LCV area was calculated (**Fig. 3C**). Contrarily, Sey1 did not promote the interaction of vacuoles harboring Δ*icmT* mutant bacteria with LDs, and overall, significantly fewer LDs associated with these vacuoles (**Fig. 3BC**). Taken together, these studies using fixed *D. discoideum* amoeba reveal that Sey1 promotes LCV-LD interactions and the Icm/Dot T4SS is required for LD accumulation on LCVs.

**Figure 3.**
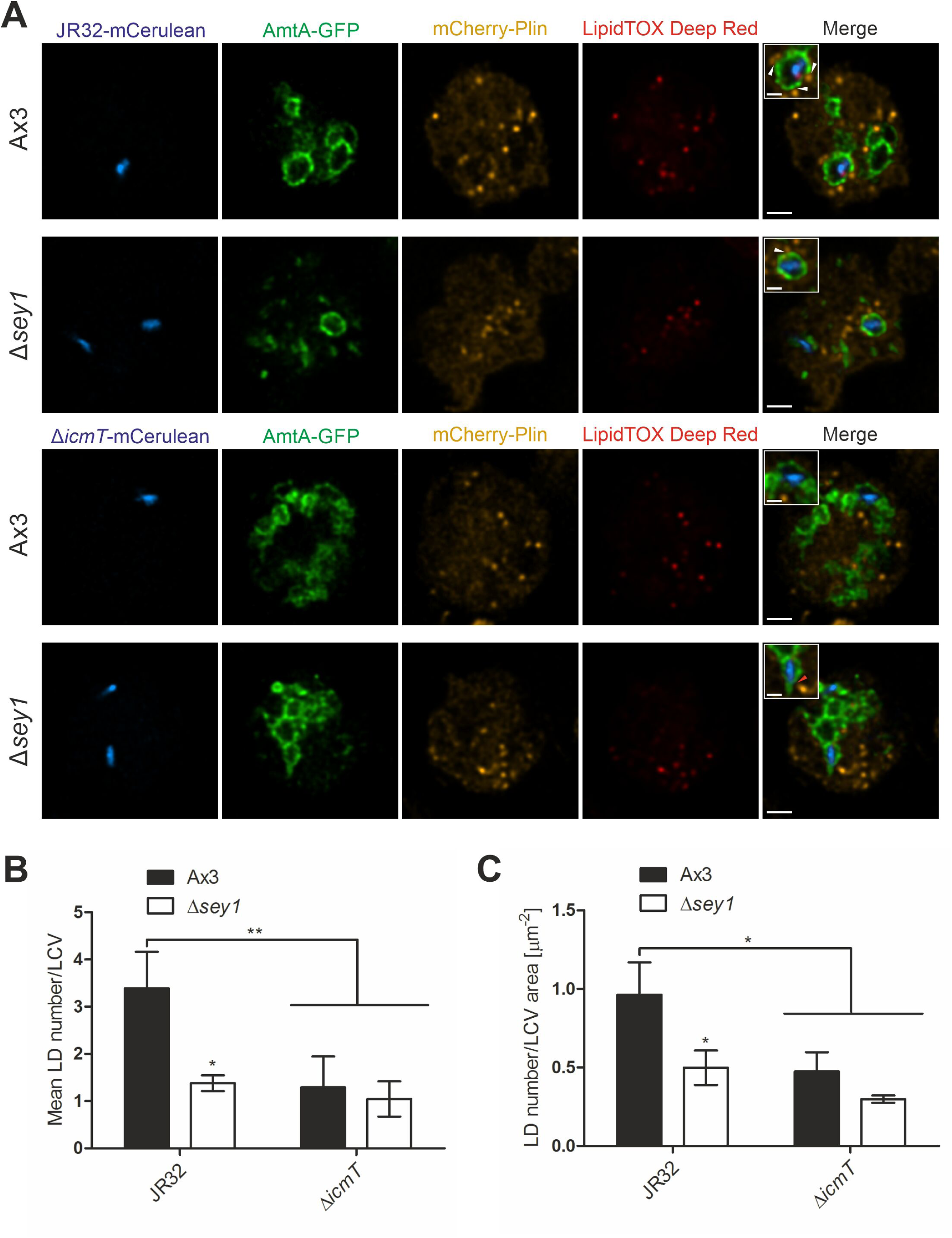
Sey1 promotes LD recruitment to LCVs in *D. discoideum*. (**A**) Representative fluorescence micrographs of *D. discoideum* Ax3 or Δ*sey1* producing AmtA-GFP (pHK121) and mCherry-Plin (pHK102), fed overnight with 200 µM sodium palmitate and infected (MOI 10, 1 h) with mCerulean-producing *L. pneumophila* JR32 (**top**) or Δ*icmT* (**bottom**) (pNP99), fixed with PFA and stained with LipidTOX Deep Red. Examples are shown for contact between LDs and the LCV membrane (**white arrowheads**) or no contact (**red arrowhead**). Scale bars: overview (2 µm), inset (1 µm). (**B**) Quantification of (A), mean LD number contacting a single LCV (n^LCVs^=30). Data represent means ± SD of three independent experiments. (\**p* < 0.05; \*\**p* < 0.01). (**C**) Quantification of (A), ratio of LD number contacting one LCV divided by the LCV area (n^LCVs^=30). Data represent means ± SD of three independent experiments. (\**p* < 0.05).

### GTP promotes LCV-LD interactions *in vitro*

Since Sey1 is a large fusion GTPase, we thought to test the nucleotide requirement of the LCV-LD interactions. To this end, we purified LCVs from *D. discoideum* Ax3 producing P4C-GFP infected with mCerulean-producing *L. pneumophila* JR32, mixed the pathogen vacuoles with purified LDs from palmitate-fed strain Ax3 producing mCherry-Plin and added 5 mM of different nucleotides (**Fig. 4A**). Using this *in vitro* reconstitution approach, the addition of GTP resulted in a ca. 2.5-fold higher number of LDs per LCV as compared to the addition of GDP, Gpp(NH)p or GTPγS (**Fig. 4B**).

**Figure 4.**
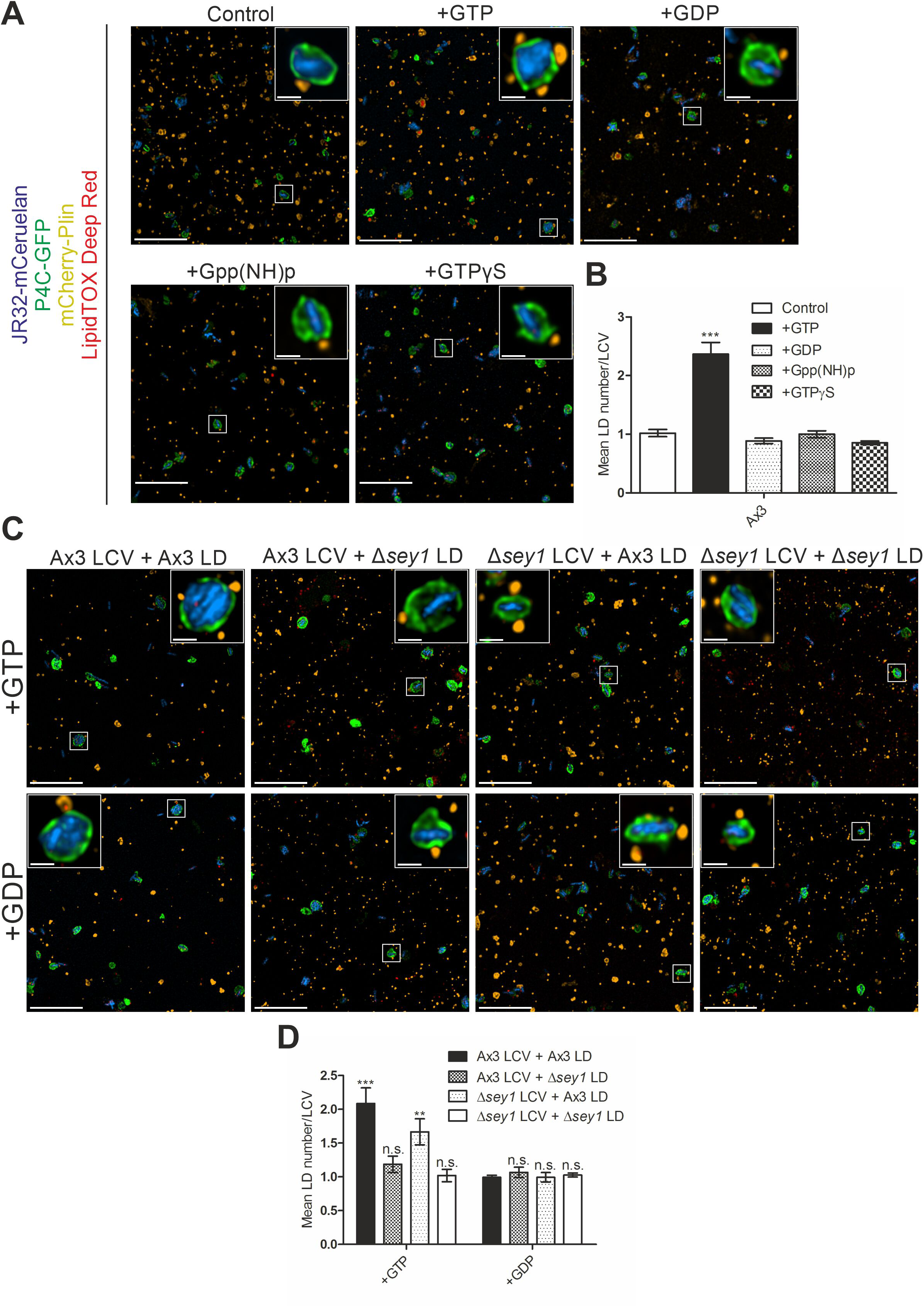
Sey1 and GTP promote LCV-LD interactions *in vitro*. (**A**) Representative fluorescence micrographs of LCVs isolated from *D. discoideum* Ax3 producing P4C-GFP (pWS034), infected (MOI 50, 1 h) with mCerulean-producing *L. pneumophila* JR32 (pNP99) and mixed with LDs from *D. discoideum* Ax3 producing mCherry-Plin (pHK102) fed overnight with 200 µM sodium palmitate. LCVs and LDs were co-incubated (1 h, 30°C) in presence of 5 mM MgCl_2_ and 5 mM GTP, GDP, Gpp(NH)p or GTPγS and fixed with PFA prior to imaging. Scale bars: overview (10 µm), inset (1 µm). (**B**) Quantification of (A), mean LD number contacting a single LCV *in vitro* (n^LCVs^=150). Data represent means ± SD of three independent experiments. (\*\*\**p* < 0.001). (**C**) Representative fluorescence micrographs of LCVs isolated from *D. discoideum* Ax3 or Δ*sey1* producing P4C-GFP (pWS034), infected (MOI 50, 1 h) with mCerulean-producing *L. pneumophila* JR32 (pNP99) and mixed with LDs from *D. discoideum* Ax3 or Δ*sey1* producing mCherry-Plin (pHK102) fed overnight with 200 µM sodium palmitate. LCVs and LDs were co-incubated (1 h, 30°C) in presence of 5 mM MgCl_2_ and 5 mM GTP (**top**) or GDP (**bottom**) and fixed with PFA prior to imaging. Scale bars: overview (10 µm), inset (1 µm). (**D**) Quantification of (C), mean LD number contacting a single LCV *in vitro* (n^LCVs^=150). Data represent means ± SD of three independent experiments. (n.s., not significant; \*\**p* < 0.01; \*\*\**p* < 0.001).

In an analogous approach, we thought to test whether Sey1 on the purified LCV or LD fraction promotes the interaction between the two compartments. We mixed LCVs purified from *D. discoideum* Ax3 or Δ*sey1* producing P4C-GFP infected with mCerulean-producing *L. pneumophila* JR32 with purified LDs from palmitate-fed strain Ax3 or Δ*sey1* producing mCherry-Plin in presence of 5 mM GTP or GDP (**Fig. 4C**). The mean number of LDs per LCV was highest for both compartments isolated from *D. discoideum* Ax3, followed by LDs purified from strain Ax3 and LCVs from Δ*sey1* mutant amoeba (**Fig. 4D**). Contrarily, the LCV-LD interactions were impaired for Δ*sey1*-derived LDs, suggesting that Sey1 regulates LD traits implicated in the interactions between the two compartments. The addition of GDP to the reconstitution assays yielded only background levels of LD/LCV ratios. In summary, *in vitro* reconstitution of the LCV-LD interactions using purified LCVs and LDs from either the *D. discoideum* Ax3 parental strain or Δ*sey1* mutant amoeba revealed that the large fusion GTPase Sey1 in the LD fraction promotes the process in a GTP-dependent manner.

### LDs shed perilipin upon LCV membrane crossing independently of Sey1 and GTP

During our analysis of LCV-LD interactions, we observed that intra-LCV LDs appeared to have lost their perilipin decoration. To analyze in more detail and quantify this observation, we used palmitate-fed *D. discoideum* Ax3 or Δ*sey1* producing P4C-GFP and mCherry-Plin. Upon staining the LDs with LipidTOX Deep Red and infection with mCerulean-producing *L. pneumophila* JR32, we quantified the portion of intra-LCV LDs without perilipin coat (**Fig. 5A**). While ca. 90% of the LDs adhering to LCVs from the cytoplasmic side were decorated with perilipin, less than 10% of the LDs in the LCV lumen were decorated with perilipin, regardless of whether the amoeba produced Sey1 or not (**Fig. 5B**). Accordingly, most LDs had shed the perilipin coat on their way from the host cell cytoplasm to the lumen of the pathogen vacuole, and this process was independent of the large fusion GTPase Sey1.

**Figure 5.**
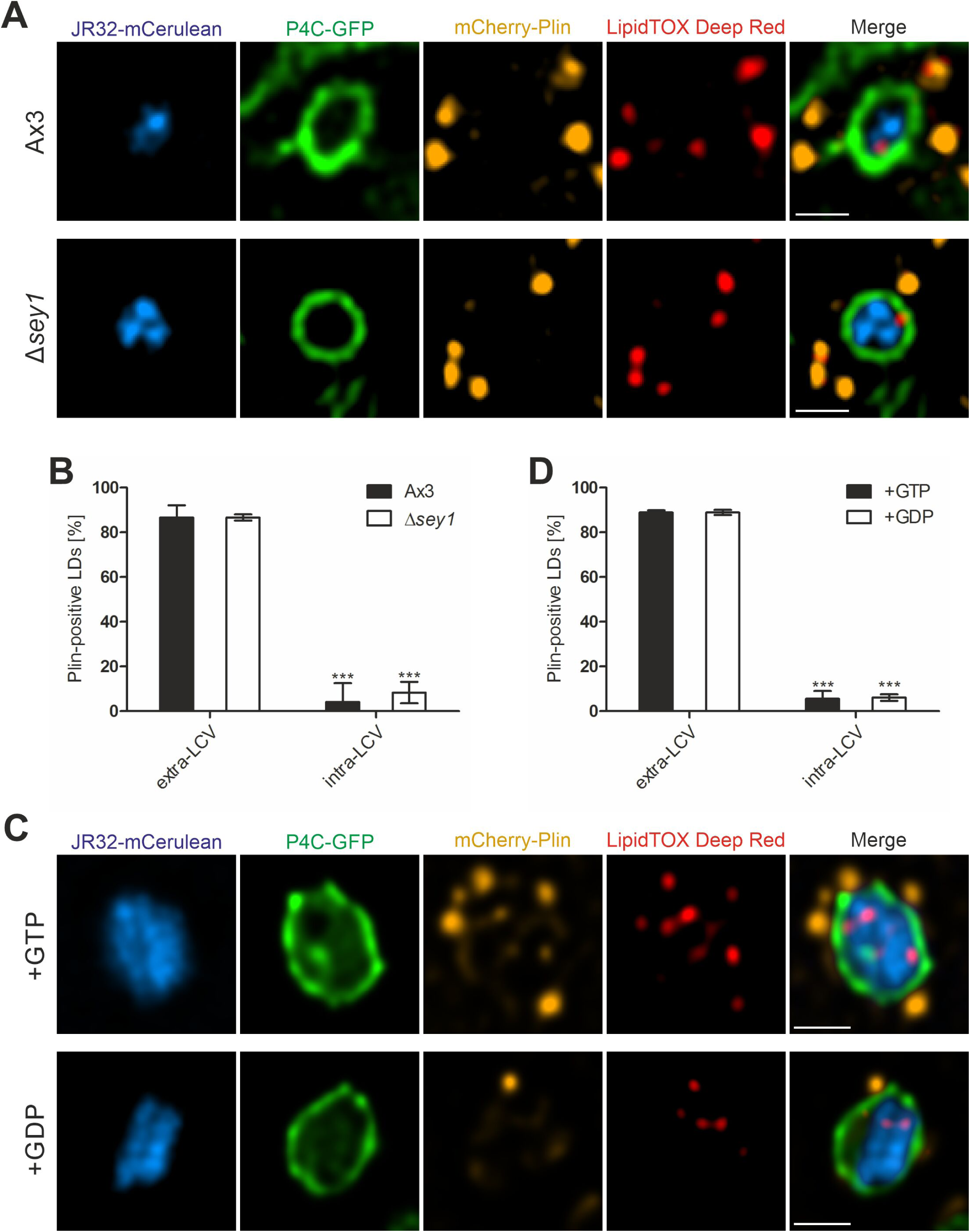
LDs shed perilipin upon LCV membrane crossing independently of Sey1 and GTP. (**A**) Representative fluorescence micrographs of *D. discoideum* Ax3 or Δ*sey1* producing P4C-GFP (pWS034) and mCherry-Plin (pHK102), fed overnight with 200 µM sodium palmitate, stained with LipidTOX Deep Red and infected (MOI 5, 1 h) with mCerulean-producing *L. pneumophila* JR32 (pNP99). Infected cells were recorded using a Leica TCS SP8. Scale bars: 1 µm. (**B**) Quantification of (A, n^LDs^ > 24); percentage of extravacuolar and intravacuolar LDs staining positive for mCherry-Plin. Data represent means ± SD of three independent experiments (\*\*\**p* < 0.001). (**C**) Representative fluorescence micrographs of LCVs isolated from *D. discoideum* Ax3 producing P4C-GFP (pWS034), infected (MOI 50, 1 h) with mCerulean-producing *L. pneumophila* JR32 (pNP99) and mixed with LDs from *D. discoideum* Ax3 producing mCherry-Plin (pHK102) fed overnight with 200 µM sodium palmitate. LCVs and LDs were co-incubated (1 h, 30°C) in presence of 5 mM MgCl_2_ and 5 mM GTP or GDP and fixed with PFA prior to imaging. Scale bars: 1 µm. (**D**) Quantification of (C, n^LDs^=180); percentage of extravacuolar and intravacuolar LDs staining positive for mCherry-Plin. Data represent means ± SD of three independent experiments (\*\*\**p* < 0.001).

To further analyze the shedding process, we mixed LCVs isolated from *D. discoideum* Ax3 producing P4C-GFP with LDs purified from palmitate-fed Ax3 producing mCherry-Plin in presence of 5 mM GTP or GDP (**Fig. 5C**). Again, ca. 90% of the LDs adhering externally to LCVs were decorated with perilipin, and less than 10% of the LDs in the LCV lumen were decorated with perilipin, regardless of whether GTP or GDP was added (**Fig. 5D**). In summary, these results indicate that upon LCV membrane crossing, LDs shed their perilipin coat in a process that does not involve Sey1 or GTP.

### Sey1 and *L. pneumophila* FadL promote intracellular replication and ^13^C_16_-palmitate catabolism

*L. pneumophila* catabolizes palmitate upon growth in broth (Häuslein et al., 2017), but it has not been analyzed whether and how the bacteria might use fatty acids during intracellular growth. Based on the observation that LDs fuse with and cross the LCV membrane concomitantly shedding their perilipin coat, LDs might deliver fatty acids to intra-vacuolar *L. pneumophila*. Moreover, fatty acids might be taken up by *L. pneumophila* through the putative outer membrane fatty acid transporter FadL. The *L. pneumophila* Philadelphia-1 genome comprises one gene that encodes a FadL homolog, *lpg1810*. The amino acid sequence of Lpg1810 is 24% identical to the sequence of *E. coli* FadL and the highly conserved NPA motif is also present (amino acids 73-75) (**Fig. S3**). The *lpg1810* gene and its genomic location are conserved among *L. pneumophila* and *L. longbeachae* strains, and the gene does not seem to be part of an operon. While genes encoding homologs of the acyl-CoA synthetase FadD, and the β-oxidation enzymes FadA, FadB and FadE are present in these *Legionella* genomes, no regulatory protein FadR homolog was identified.

We first tested whether *fadL* (*lpg1810*) is expressed and how expression is controlled. To this end, we constructed a P*_fadL_*-*gfp* transcriptional fusion and assessed GFP production in *L. pneumophila* JR32 and Δ*lqsA*, Δ*lqsR,* or Δ*lvbR* mutant strains lacking components of the Lqs quorum sensing system or the pleiotropic transcription factor LvbR, respectively (**Fig. S4AB**). Compared to the parental strain JR32, P*_fadL_*-*gfp* was expressed less strongly and with some delay in the Δ*lvbR* mutant but similarly in the Δ*lqsA* and Δ*lqsR* mutants. These results indicate that the transcription factor LvbR but not the Lqs quorum sensing system regulates the expression of *fadL* under the conditions tested.

To assess the role of *fadL* for the growth of *L. pneumophila*, we constructed a Δ*fadL* deletion mutant strain by double homologous recombination. The Δ*fadL* mutant strain grew like the parental strain JR32 in AYE broth and MDM minimal medium (**Fig. S4C**). However, the Δ*fadL* mutant strain was impaired for intracellular growth in *D. discoideum*, and the growth defect was complemented by inserting the *fadL* gene back into the *L. pneumophila* genome (**Fig. 6A**). In agreement with a role of *fadL* for intracellular growth of *L. pneumoph*ila, the P*_fadL_* reporter construct was expressed in *D. discoideum* throughout an infection (**Fig. S5**).

**Figure 6.**
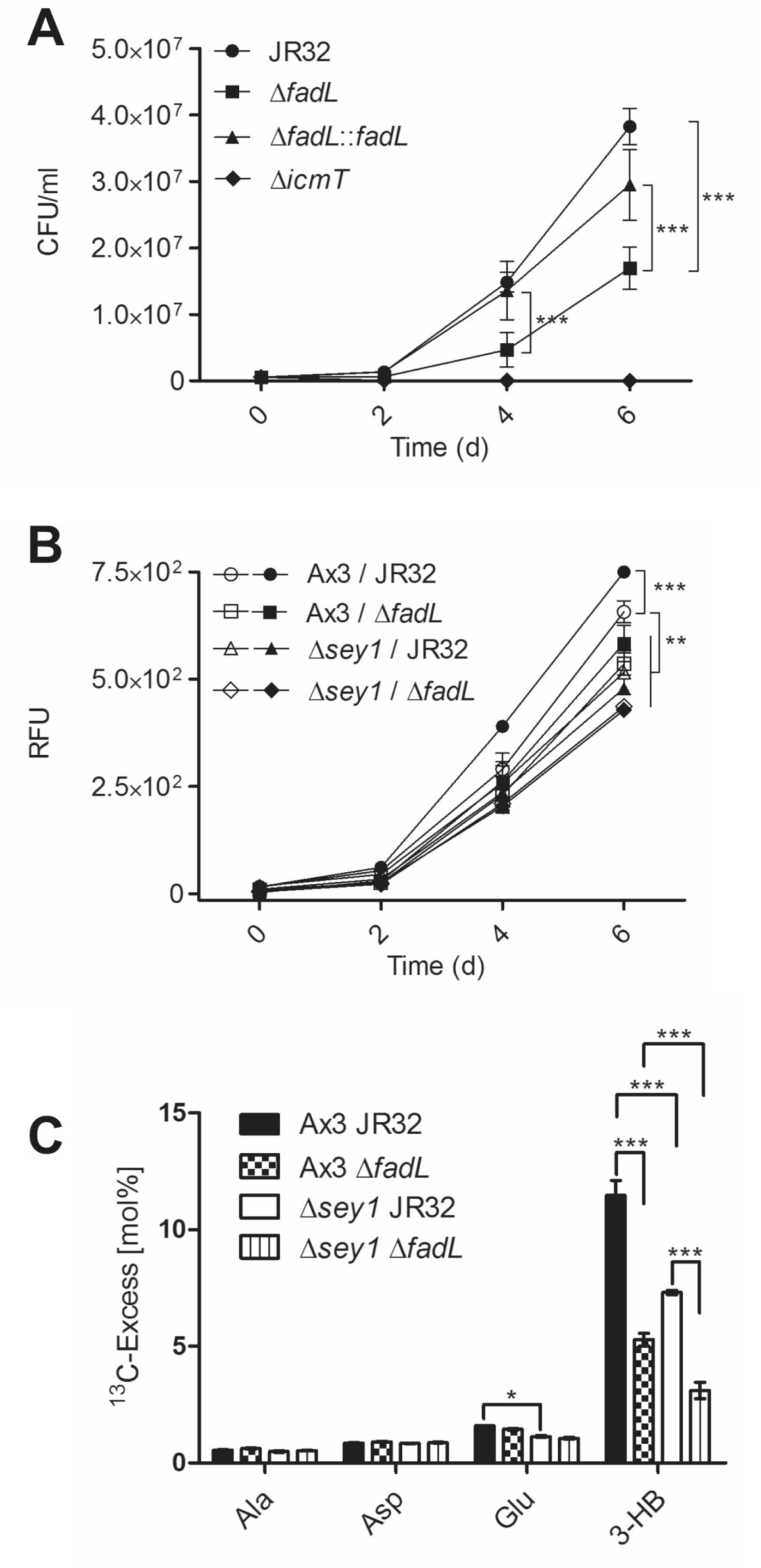
*D. discoideum* Sey1 and *L. pneumophila* FadL promote intracellular pathogen replication and ^13^C_16_-palmitate catabolism. (**A**) *D. discoideum* Ax3 were infected (MOI 1) with *L. pneumophila* JR32, Δ*icmT*, Δ*fadL* or Δ*fadL*::*fadL* (chromosomal integration of *fadL* in Δ*fadL*), and intracellular replication was assessed by CFU for 6 days. Data represent means ± SD of three independent experiments in technical triplicates (\*\*\**p* < 0.001). (**B**) *D. discoideum* Ax3 or Δ*sey1* were left untreated (**empty symbols**) or fed overnight with 200 µM sodium palmitate (**filled symbols**) and were infected (MOI 1) with GFP-producing *L. pneumophila* JR32 or Δ*fadL* (pNT28) for 6 days. Intracellular replication of the bacteria was assessed by quantification of relative fluorescence units (RFUs). Data represent means ± SD of three independent experiments in technical hextuplicates (\*\**p* < 0.01; \*\*\**p* < 0.001). (**C**) *D. discoideum* Ax3 or Δ*sey1* producing calnexin-GFP (CnxA-GFP, pAW016) were infected (MOI 50, 1 h) with mCerulean-producing *L. pneumophila* JR32 or Δ*fadL* (pNP99) and washed to remove extracellular bacteria. At 5 h p.i., 200 µM [U-^13^C_16_]palmitate was added to the infected amoeba for 10 h. The infected cells were lysed and centrifuged to separate bacteria from cell debris. ^13^C-excess (mol%) in key metabolites of the *L. pneumophila* fraction was analyzed by GC/tandem MS. Data show means ± SD of technical triplicates (\*\*\**p* < 0.001) and are representative for two independent experiments.

Next, we tested the effects of overnight palmitate feeding of *D. discoideum* Ax3 or Δ*sey1* on intracellular growth of *L. pneumophila* JR32 or Δ*fadL*. Feeding with palmitate augmented the growth of *L. pneumophila* JR32 in *D. discoideum* Ax3 but did not significantly affect the growth of the Δ*fadL* strain in either *D. discoideum* Ax3 or Δ*sey1* (**Fig. 6B**). These results revealed that the host large fusion GTPase Sey1 as well as the putative *L. pneumophila* fatty acid transporter FadL are required for intracellular growth promotion by palmitate and presumably the catabolism of this carbon and energy source.

Finally, to test whether Sey1 and FadL indeed promote the intracellular catabolism of palmitate, we used the isotopologue profiling method. With this approach, the intracellular catabolism by *L. pneumophila* of ^13^C-labeled carbon sources via ^13^C_2_-acetyl-CoA and incorporation of label into the storage compound ^13^C-poly-3-hydroxybutyrate (PHB) is followed by MS (Eylert et al., 2010; Schunder et al., 2014; Häuslein et al., 2016). To test intracellular catabolism of palmitate, calnexin-GFP-producing *D. discoideum* Ax3 or Δ*sey1* were infected with mCerulean-producing *L. pneumophila* JR32 or Δ*fadL*, treated with 200 µM [U-^13^C_16_]palmitate for 10 h, lysed and fractionated. ^13^C-Excess (mol%) in key metabolites of the *L. pneumophila* fraction was analyzed by gas chromatography/MS (**Fig. 6C**). This approach revealed a minor enrichment of ^13^C label in the amino acids Ala, Asp and Glu; in Glu, the enrichment was less efficient in absence of Sey1. A major enrichment of ^13^C-label occurred in the PHB hydrolysis product 3-hydroxybutyrate (3-HB), and interestingly, the enrichment was significantly lower in absence of Sey1 or FadL, and lowest in absence of both, Sey1 and FadL. In summary, these studies revealed that the host large fusion GTPase Sey1 as well as the outer membrane fatty acid transporter FadL are implicated in intracellular palmitate catabolism by *L. pneumophila*.

## Discussion

In this study we assessed the role of LDs, Sey1 and FadL for intracellular growth of *L. pneumophila*. We reveal that LCVs intimately interact with palmitate-induced and Sey1-containing LDs in *D. discoideum* (**Fig. 1**). Moreover, using dually fluorescence-labeled amoeba, we demonstrated *in vivo* by live-cell microscopy (**Fig. 2**) and fixed samples (**Fig. 3**), or *in vitro* using reconstituted purified LCVs and LDs (**Fig. 4**) that Sey1 and GTP promote LCV-LD interactions. Palmitate was catabolized and promoted the intracellular replication of wild-type *L. pneumophila* in *D. discoideum* Ax3, but not of Δ*fadL* mutant bacteria in Δ*sey1* mutant amoeba (**Fig. 6**). Taken together, our results indicate that Sey1-dependent recruitment of LDs, LCV-LD interactions, LD transfer to the LCV lumen and FadL-dependent catabolism of fatty acids promote intracellular growth of *L. pneumophila* (**Fig. 7**).

**Figure 7.**
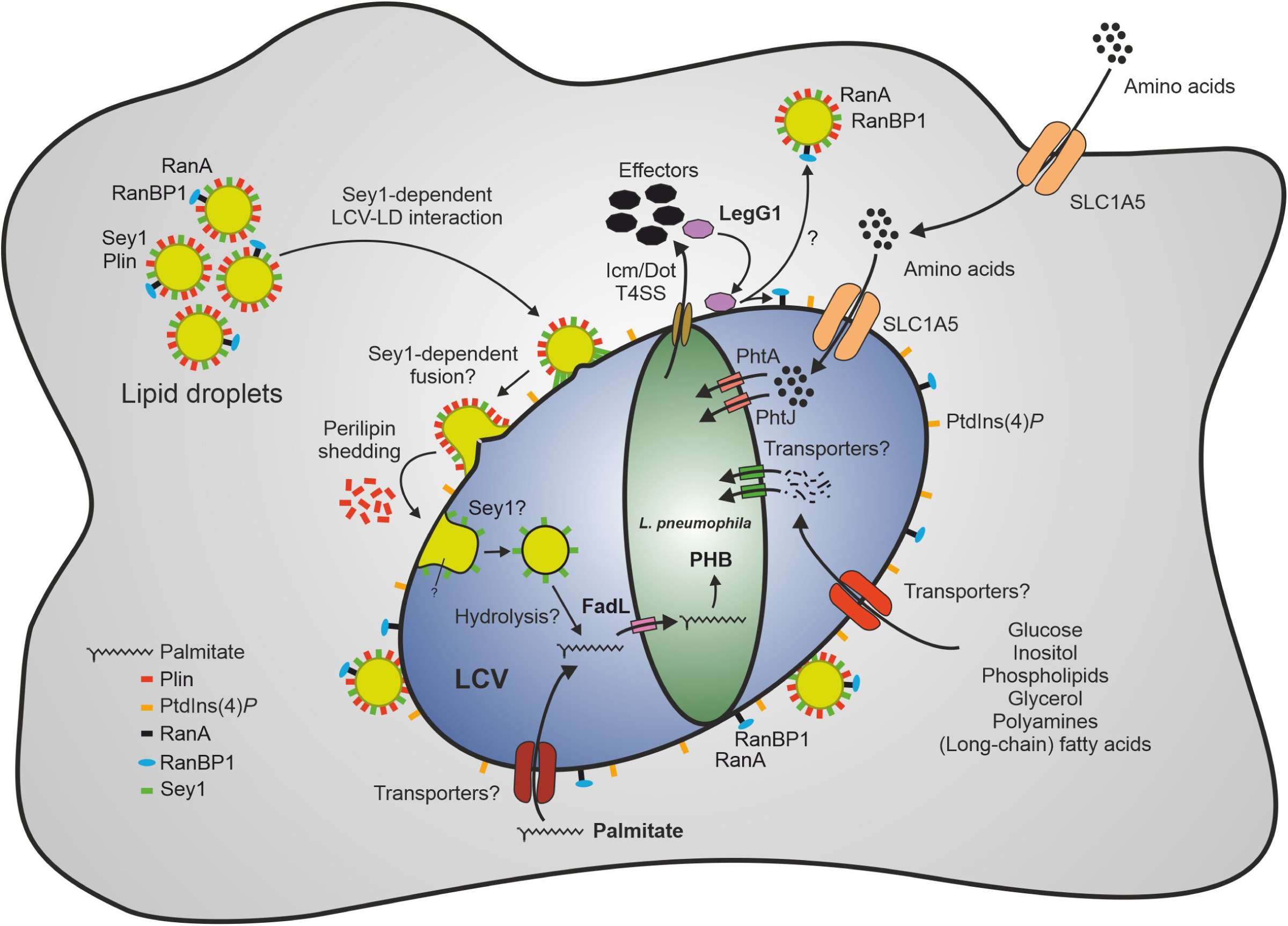
Sey1-, LegG1- and FadL-dependent catabolism of palmitate by *L. pneumophila* through lipid droplets. The large fusion GTPase Sey1 and GTP as well as the *L. pneumophila* RCC1 repeat effector LegG1 promote the recruitment of host LDs to LCVs. LegG1 activates the small GTPase Ran, leading to accumulation of RanBP1 and microtubule stabilization. Upon intimate contact of LDs with LCVs, perilipin is shed, and LDs cross the membrane to reach the LCV lumen, where LD constituents (e.g., triacylglycerols) are likely hydrolyzed, and free fatty acids are taken up by *L. pneumophila*. Palmitate is transported by FadL inside the bacteria and metabolized to acetyl-CoA, which is further aerobically catabolized in the tricarboxylic acid cycle to CO_2_ and/or anabolized to the storage compound polyhydroxybutyrate (PHB). Amino acids are transported by the host transporter SLC1A5 and the *L. pneumophila* transporters PhtA/PhtJ, respectively. Additional trans-membrane transporters for lipids (long-chain fatty acids, phospholipids), sugars (glucose, inositol), alcohols (glycerol), polyamines, and amino acids are likely present in the plasma membrane, LCV membrane and *L. pneumophila* membrane, respectively. LD, lipid droplet; LCV, *Legionella*-containing vacuole; PHB, polyhydroxybutyrate; Plin, perilipin; RanBP1, Ran-binding protein 1.

The LCV-LD interactions are controlled by the *L. pneumophila* Icm/Dot T4SS (**Fig. 3**), and the Icm/Dot substrate LegG1 (**Fig. 2**). LegG1 belongs to a family of RCC1 repeat domain-containing *L. pneumophila* effectors, which activate the small GTPase RanA at different sites in the cell (LCV, plasma membrane) and consequently stabilize non-centrosomal microtubules (Rothmeier et al., 2013; Swart et al., 2020a; Swart et al., 2020b). In addition to RanA, the Ran-binding protein RanBP1 was identified on LDs (**Fig. S2A**). Since RanBP1 only binds to activated, GTP-bound RanA, the small GTPase is likely activated on LDs. Accordingly, LegG1 might not only activate RanA on LCVs but also on LDs, leading to a stabilization of microtubules, along which the LCV-LD interactions occur.

Among the more than 330 different *L. pneumophila* effectors, others might (directly) target LDs. An attractive candidate is RalF, which acts as an Arf1 guanine nucleotide exchange factor (GEF), thus activating the small GTPase Arf1 on LCVs (Nagai et al., 2002). Activated, GTP-bound Arf1 regulates through perilipin-2 the phospholipid monolayer that stabilizes LDs and controls the cellular availability of neutral lipids for metabolic purposes (Pauloin et al., 2016). Moreover, upon activation, Arf1 and the coat complex COPI form LD-ER membrane bridges for targeting catabolic enzymes (Wilfling et al., 2014). Hence, by acting as an Arf1 GEF, the *L. pneumophila* effector RalF might regulate LD-dependent processes during *L. pneumophila* infection.

In addition to *L. pneumophila*, several intracellular pathogens accumulate LDs on their pathogen vacuoles to get access to and metabolize fatty acids (Walpole et al., 2018; Bosch et al., 2021; Brink et al., 2021). Pathogenic *Mycobacterium* species, such as *M. tuberculosis* and *M. leprae*, trigger LD formation during infection, to exploit lipids derived from these organelles as carbon and energy source. *M. tuberculosis* actively evokes LDs in macrophages, thus promoting the “foamy” appearance of these host cells, to employ lipids as primary energy source in the “dormant” state during chronic infection (Singh et al., 2012; Mehrotra et al., 2014). Moreover, LDs accumulate around *Mycobacterium*-containing vacuoles (MCVs) harboring *M. marinum* in *D. discoideum* (Barisch et al., 2015; Barisch and Soldati, 2017a, b). These LDs cross to the lumen of the MCV and form “intracytosolic lipid inclusions” (ILIs) within the bacteria. However, during the *D. discoideum*-*M. marinum* encounter, ILI formation and lipid catabolism apparently do not lead to dormancy of the bacteria.

The obligate intracellular pathogen *Chlamydia trachomatis* forms a pathogen vacuole termed the inclusion (Fields and Hackstadt, 2002). The inclusion accumulates various host lipids, such as sphingomyelin (Hackstadt et al., 1996; Elwell et al., 2011) and cholesterol (Carabeo et al., 2003), and sphingomyelin promotes intracellular growth (van Ooij et al., 2000; Elwell and Engel, 2012). Intriguingly, the *C. trachomatis* inclusions also capture LDs (Kumar et al., 2006), which transverse across the inclusion membrane to the lumen of the pathogen compartment (Cocchiaro et al., 2008). LDs seem to be a source of lipids for *C. trachomatis*, but they are not essential as a delivery carrier of fatty acids (Sharma et al., 2018). Fatty acids are activated by host long chain fatty acid Acyl-CoA synthases, which are recruited to the inclusion (Recuero-Checa et al., 2016).

*L. pneumophila* FadL promotes the catabolism of palmitate (**Fig. 6**), but overall, the degradation of lipids by *Legionella* is not well understood. Analogously to *M. marinum* (Barisch and Soldati, 2017b), *L. pneumophila* might hydrolyse triacylglycerols; however, rather than building up ILIs, *L. pneumophila* accumulates the storage compound PHB (**Fig. 6**). *L. pneumophila* produces at least 19 phospholipases (15 phospholipases A, three phospholipases C, and one phospholipase D), which act as virulence factors during intracellular replication (Kuhle and Flieger, 2013), and some of which might metabolize lipids stored in LDs. While some of the lipases are T2SS substrates and thus are presumably secreted to the LCV lumen, others are Icm/Dot T4SS substrates and translocated to the host cytoplasm. Type II-secreted phospholipases comprise phospholipases A, i.e., the GDSL lipase family members PlaA and PlaC, or PlaB and PlaD (Bender et al., 2009; Lang et al., 2017), as well as the zinc metallophospholipases C, i.e., PlcA and PlcB (Aurass et al., 2013). PlcC (CegC1/Lpg0012) is a type IV (Icm/Dot)-translocated zinc metallophospholipases C (Aurass et al., 2013). Alternatively, or additionally, some of these phospholipases might be involved in degrading the limiting LCV membrane to escape the pathogen vacuole and the host cell. In agreement with this notion, *L. pneumophila* triple mutant strain lacking phospholipases A or C were impaired for LCV rupture in *D. discoideum*, indicating that phospholipases A and C are indeed involved in membrane lysis (Striednig et al., 2021). Further studies will address the role of these phospholipases for *L. pneumophila* LD utilization and lipid metabolism.

Mechanistically, the fusion of LDs with distinct pathogen vacuoles is poorly understood (**Fig. 1A**). In particular, it is unknown how LDs tether and cluster on pathogen vacuoles and how they cross the pathogen vacuole membrane and translocate inside the pathogen vacuole. Given the role of Sey1 for the recruitment, accumulation and contact time of LDs on LCVs, the large fusion GTPase might also play a role in later steps of LD-LCV fusion and membrane crossing. Sey1 was identified by proteomics on LDs purified from the *D. discoideum* Ax3 parental strain, but not from the Δ*sey1* mutant strain (**Fig. 1B**). Accordingly, Sey1 might directly participate in the LD-LCV fusion process. However, we cannot rule out that the ER-residing fusion GTPase Sey1 affects the formation and composition of ER-derived LDs, and thus, indirectly affects LD-LCV fusion. Since Sey1 does not seem to significantly affect the number and size of LDs in *D. discoideum* (**Fig. S1BC**), an indirect role of Sey1 for LD-LCV interactions seems less likely.

The LD protein perilipin was shed upon uptake of LDs into LCVs (**Fig. 5**). Perilipin reversibly interacts with the LD surface via amphipathic helices (Kory et al., 2016), and therefore, it might be rather easily shed and distributed to the cytoplasm upon LD translocation to the LCV lumen. The mechanism and biological function of perilipin shedding upon pathogen vacuole transfer of the LDs is unknown. Future investigations using the LCV-LD model system for *in vivo* and *in vitro* studies by confocal microscopy and reconstitution approaches, respectively, will provide further insights on the underlying cell biological and infection biological processes.

## Material and methods

### Bacteria, cells, and growth conditions

Bacterial strains and cell lines used are listed in **Table S2**. *L. pneumophila* strains were grown for 3 days on charcoal yeast extract (CYE) agar plates, buffered with *N*-(2-acetamido)-2-aminoethane sulfonic acid (ACES) at 37°C. Liquid cultures in ACES yeast extract (AYE) medium were inoculated at an OD_600_ of 0.1 and grown at 37°C for 21 h to an early stationary phase (2×10^9^ bacteria/ml). Chloramphenicol (Cam; 5 μg/ml) was added when required.

Growth of *L. pneumophila* strains was assessed in 3 ml AYE or 3 ml minimal defined medium (MDM) at an initial OD_600_ of 0.1 and incubated on a rotating wheel (80 rpm, 37°C) for 18 h or 28 h, respectively. The cultures were then diluted in the respective medium to an OD_600_ of 0.2 in a 96-well plate (200 μl/well) and incubated at 37°C while orbitally shaking. The OD_600_ was measured in triplicates using a microtiter plate reader to assess growth in AYE (Synergy H1 Hybrid Reader, BioTek) or MDM (Cytation 5 Cell Imaging Multi-Mode Reader; BioTek).

For GFP reporter assays, bacterial overnight cultures were diluted to an initial OD_600_ of 0.2. In a black 96-well plate, 200 μl diluted over-night culture per well was incubated at 37°C while orbitally shaking. Bacterial growth and GFP production were monitored in triplicates by measuring the OD_600_ and fluorescence (excitation, 485 nm; emission, 528 nm; gain, 50) using a microtiter plate reader (Cytation 5 Cell Imaging Multi-Mode Reader, BioTek). Values are expressed as relative fluorescence units (RFU) or OD_600_.

*D. discoideum* strains were grown at 23°C in HL-5 (ForMedium) medium. Cells were maintained every 2-3 days by rinsing with fresh HL-5 and by transferring 1-2% of the volume to a new T75 flask containing 10 mL medium. Cells were strictly maintained below 90% confluence. Transformation of axenic *D. discoideum* amoeba was performed as previously described (Weber and Hilbi, 2014; Weber et al., 2014). Geneticin (G418, 20 µg/mL) and hygromycin (50 µg/mL) were added when required.

### Molecular cloning

The plasmids used in this study are listed in **Table S2**. Cloning was performed according to standard protocols and plasmids were isolated using the NucleoSpin Plasmid kit (Macherey-Nagel). DNA fragments were amplified using Phusion High Fidelity DNA polymerase (NEB) and the oligonucleotides listed in **Table S3**. FastDigest restriction enzymes (Thermo-Fisher) were used for plasmid digestion and Gibson assembly was performed using the NEBuilder HiFi DNA assembly kit (NEB). All constructs were verified by DNA sequencing.

To construct the plasmid pLS187, the gene region of *ranBP1* (P*_ranBP1_*) was amplified from pSU26 (Rothmeier et al., 2013) using the primer pair oLS296/oLS297. The PCR product was purified as described above and cloned into pDM323 (Veltman et al., 2009), previously digested with *Bgl*II and *Spe*I. To construct pLS221 and pLS222, the gene region of *ranA* (P*_ranA_*) and *ranBP1* (P*_ranBP1_*) was amplified from pSU17 and pLS187, respectively, using the primer pairs oLS333/oLS334 and oLS296/oLS297. The PCR products were purified as described above and cloned into the plasmids pDM1044 (Barisch et al., 2015) or pDM329 (Veltman et al., 2009), respectively, digested with *Bgl*II and *Spe*I.

To construct the GFP reporter plasmid pPS003, the promoter region of *fadL* (P*_fadL_*) was amplified from genomic DNA of *L. pneumophila* using the primers oPS013 and oPS015. The PCR product was purified using the NucleoSpin Gel and PCR Clean-up kit (Macherey-Nagel) and cloned into the plasmid pCM009 (Schell et al., 2016), which was previously digested with *Sac*I and *Xba*I to remove P*_flaA_*. Thus, P*_flaA-_gfp* was replaced by P*_fadL_*-*gfp* in the final construct.

### Construction of the *L. pneumophila* Δ*fadL* mutant

The Δ*fadL* mutant was generated by double homologous recombination as described (Tiaden et al., 2007), replacing the *fadL* gene by a kanamycin (Kan) resistance cassette. The primers oPS001 and oPS002 were used to amplify from *L. pneumophila* genomic DNA the upstream flanking region of *fadL*, oPS005 and oPS006 to amplify the downstream flanking region of *fadL* (939 bp each). The Kan resistance cassette was amplified using the primers oPS003 and oPS004, and the plasmid pUC4K (Amersham) as a template. The three amplified fragments were cloned into *Bam*HI-digested pUC19 in a 3-way ligation and amplified in one piece using the primers oPS010 and oPS011. To yield the allelic exchange vector pPS002, the amplified sequence was cloned into the *Bam*HI-digested suicide vector pLAW344 (Wiater et al., 1994), which allows counter-selection with the *sacB* gene. *L. pneumophila* JR32 was transformed with pPS002 by electroporation and grown on CYE plates supplemented with 5 µg/ml Cam. Individual clones were picked, grown in AYE medium overnight and plated on CYE plates supplemented with 20 µg/ml Kan and 20 mg/ml sucrose, or 5 µg/ml Cam. After incubation at 37°C for 2 days, clones growing on the CYE/Kan/sucrose plate were picked and purified by dilution streaking. This selection process was repeated until growth on CYE/Cam plates was no longer observed. Double-cross-over events and thus correct insertion of the Kan resistance cassette in the genome of the deletion mutant were confirmed by PCR and sequencing.

For complementation of the intracellular replication phenotype of Δ*fadL*, the *fadL* gene was re-introduced into the genome of the Δ*fadL* mutant strain by co-integration of the suicide plasmid pPS013. This plasmid was constructed by amplification of *fadL* together with its 3’ and 5’ flanking regions from *L. pneumophila* genomic DNA using the primers oPS011 and oPS045 and cloning into *Bam*HI-digested pPS002. Transformants were plated on CYE plates supplemented with 5 µg/ml Cam to select for uptake and integration of the pPS013.

### Visualization and quantification of *fadL* expression

To assess *fadL* (*lpg1810*) expression in planktonic *L. pneumophila*, liquid cultures of strains harboring pPS003 were prepared as described above and incubated on a rotating wheel for the time indicated. Subsequently, 100 μl liquid culture was collected and centrifuged (5 min, 2000 *g*, room temperature, RT). The bacteria were fixed in 4% paraformaldehyde (PFA, 1 h, RT), stained with 1 µg/ml DAPI in DPBS (1 h, RT), and washed twice with DPBS. For microscopy, the bacteria were pelleted, resuspended in 30 µl DPBS and embedded in 18-well ibidi dishes in 0.5% agarose in DPBS. For flow cytometry, pellets were resuspended in 500 µl DPBS.

To assess *fadL* expression in intracellular *L. pneumophila*, *D. discoideum* were infected with *L. pneumophila* JR32 harboring pPS003. Amoebae were counted in a Neubauer improved counting chamber, 0.1 mm depth (Marienfeld-Superior) and 1×10^6^ *D. discoideum* per well were seeded one day before infection into 6-well plates in 2 ml HL-5. *L. pneumophila* overnight cultures were prepared as described above and grown on a rotating wheel at 37°C for 21-22 h to early stationary phase (OD_600_ ca. 5.0, ∼2×10^9^ bacteria/ml). The day of infection, *L. pneumophila* cultures were checked for motility and filamentation, and OD_600_ was measured to calculate the bacterial concentration. *D. discoideum* cells were infected (MOI 5), centrifuged (10 min, 450 *g*, RT), and incubated at 25°C. After 1.5-2 h, *D. discoideum* cells were washed four times with HL-5, and incubated further at 25°C. For the 30 min p.i. time point, amoebae were washed after 30 min and immediately processed further. Infected amoebae were collected into 2 ml Eppendorf tubes 30 min, 6 h, 24 h, 32 h, or 48 h p.i., and centrifuged (5 min, 2000 *g*, RT) to remove the medium. For microscopy, the cells were fixed in 4% PFA (1 h, RT), washed with DPBS, and permeabilized with ice-cold methanol for 10 min. Subsequently, cells were stained with 1 µg/ml DAPI (1 h, RT), washed twice with DPBS, resuspended in 30 µl DPBS, and embedded in 0.5% agarose in an 18-well ibidi dish.

For flow cytometry, cells were lysed in lysis buffer (150 mM NaCl, 0.1% Triton X-100) for 15 min at RT and centrifuged (5 min, 2000 *g*, RT). Supernatant was removed and isolated intracellular bacteria were fixed in 4% PFA (1 h, RT). The samples were stained with 1 µg/ml DAPI (1 h, RT), washed twice with DPBS, and resuspended in 500 µl DPBS.

### Intracellular *L. pneumophila* replication

Intracellular replication of *L. pneumophila* JR32, Δ*icmT* and Δ*fadL* in *D. discoideum* amoebae was analyzed by colony forming units (CFUs) as well as by increase in relative fluorescence units (RFUs). To determine CFUs, *D. discoideum* Ax3 or Δ*sey1* amoebae were seeded at a density of 1×10^5^ cells/ml in cell culture-treated 96-wells plates (VWR) and cultivated at 23°C in HL-5 medium. Afterwards, the amoebae were infected (MOI 1) with early stationary phase *L. pneumophila* JR32, Δ*icmT*, Δ*fadL*, or Δ*fadL*::*fadL* (chromosomal integration of *fadL* in Δ*fadL*) diluted in MB medium (Solomon and Isberg, 2000), centrifuged (450 *g*, 10 min, RT) and incubated at 25°C for the time indicated (96-wells plate was kept moist by addition of ddH_2_O in surrounding wells). The cells were lysed with 0.8% saponin (Sigma-Aldrich) for 10 min at RT, and dilutions were plated on CYE/Cam agar plates and incubated at 37°C for 3 days. CFUs were assessed every 2 days using an automated colony counter (CounterMat Flash 4000, IUL Instruments, CounterMat software), and the number of CFUs (per ml) was calculated.

To determine increase in fluorescence derived from intracellular replication of GFP-producing *L. pneumophila*, *D. discoideum* Ax3 or Δ*sey1* were seeded at a density of 1×10^5^ cells/ml in cell culture-treated 96-wells plates (VWR) and cultivated overnight at 23°C in HL-5 medium supplemented with 200 µM sodium palmitate. A range of different concentrations of sodium palmitate (100-800 µM) were previously tested, and 200 µM sodium palmitate appeared to be the most effective in increasing intracellular replication of *L. pneumophila* (**Fig. S1A**). The cells were infected (MOI 1) with early stationary phase GFP-producing *L. pneumophila* JR32 or Δ*fadL* diluted in MB medium, centrifuged (450 *g*, 10 min, RT) and incubated for 1 h at 25°C. Afterwards, the infected amoebae were incubated at 25°C for the time indicated (96-wells plate was kept moist by addition of ddH_2_O in surrounding wells). Increase in GFP fluorescence was assessed every 2 days using a microtiter plate reader (Synergy H1, Biotek).

### Flow cytometry

Intracellular bacteria recovered from infected amoebae or strains grown in liquid culture were analyzed with an LSRFortessa II (BD Biosciences). Gating was performed as follows: Bacterial cells were identified by forward scatter (FSC) versus side scatter (SSC) gating. The threshold for FSC and SSC was set to 200, and 10’000 events were acquired per sample. The bacteria were further examined for their DAPI (450 nm) and GFP (488 nm) fluorescence. DAPI-stained JR32 and JR32 harboring pNT28 for constitutive GFP production were used as references for gating of the GFP-positive subpopulation in the samples. Thus, the percentage of GFP-positive bacteria was calculated. Data processing was performed using the FlowJo software.

### In vitro reconstitution assays

LCVs from *D. discoideum* Ax3 and Δ*sey1* mutant amoebae were purified as previously described (Urwyler et al., 2010). Briefly, *D. discoideum* Ax3 or Δ*sey1* producing P4C-GFP (pWS034) were seeded in three T75 flasks per sample one day prior to experiment to reach 80% confluency. The amoebae were infected (MOI 50, 1 h) with *L. pneumophila* JR32 producing mCerulean (pNP99) grown to stationary phase (21 h liquid culture). Subsequently, the cells were washed with SorC buffer (2 mM Na_2_HPO_4_, 15 mM KH_2_PO_4_, 50 µM CaCl_2_ × 2H_2_O, pH 6.0) and scraped in homogenization buffer (20 mM HEPES, 250 mM sucrose, 0.5 mM EGTA, pH 7.2) containing a protease inhibitor cocktail tablet (Roche) (Derre and Isberg, 2004). Cells were homogenized using a ball homogenizer (Isobiotec) with an exclusion size of 8 µm and incubated with an anti-SidC antibody followed by a secondary anti-rabbit antibody coupled to magnetic beads. The LCVs were separated using magnetic columns and further purified by density gradient centrifugation as described (Hoffmann et al., 2013).

LDs from *D. discoideum* Ax3 and Δ*sey1* mutant amoebae were purified using an updated protocol as previously published (Du et al., 2013). *D. discoideum* Ax3 or Δ*sey1* producing mCherry-Plin (pHK102) were seeded in 3 x T150 flasks per sample one day prior to experiment to reach 80% confluency and stimulated overnight (16 h) with 200 µM sodium palmitate to induce LDs formation. The next day, the cells were recovered by scraping, washed twice with SorC buffer and once in STKM buffer (50 mM Tris, pH 7.6, 25 mM KCl, 5 mM MgCl_2_ and 0.25 M sucrose) supplemented with a protease inhibitor cocktail tablet (Roche). Successively, cells were resuspended in 1 ml STKM buffer (supplemented with protease inhibitor cocktail tablet) and homogenized by 9-10 passages through a ball homogenizer (Isobiotec) with an exclusion size of 8 µm. Cell homogenate was then centrifuged (1000 *g*, 30 min, 4°C) and the supernatant containing LDs was separated from the nuclei pellet. The post-nuclear supernatant was adjusted to 0.8 M sucrose and loaded in the middle of a step gradient ranging from 0.1 to 1.8 M sucrose in STKM buffer and centrifuged at 50’000 g for 1 h at 4°C in a TLA-120.1 fixed-angle rotor (Beckman Coulter). LDs formed a white cushion on top of the tube and were collected by aspiration of the upper fraction of the gradient (125 µl).

Isolated LDs and LCVs were combined *in vitro* to reconstitute and biochemically analyze LCV-LD interaction. Isolated LCVs (300 µl from gradient) were diluted with 1 ml homogenization buffer with protease inhibitors, added on top of sterile poly-L-lysine (Sigma) coated coverslips in a 24-well plate and centrifuged at 600 *g* for 10 min for 4°C. Supernatant was aspirated and isolated LDs (100 µl from gradient), together with 100 µl STKM with protease inhibitors, 5mM MgCl_2_ and 5mM GTP/GDP/Gpp(NH)p/GTPγS (Sigma) were appropriately mixed and added on top of the LCV-coated coverslips. Reconstitution plate was then centrifuged (600 *g*, 10 min, 4°C) and incubated for 1 h at 30°C. Following incubation and supernatant aspiration, *in vitro* LCV-LD were fixed with 2% PFA for 1 h at room temperature, washed twice with SorC, stained with LipidTOX Deep Red (1:200 in SorC; Thermo-Fisher) for 30 min at room temperature in the dark, and finally mounted on microscopy slides.

### Confocal microscopy of bacteria, infected cells, and LCV-LD interactions *in vitro*

For infection assays for confocal microscopy, *D. discoideum* strains producing the desired fluorescent probes were harvested from approximately 80% confluent cultures, seeded at 1×10^5^ cells/mL in 6-well plates (Corning) or 8-well µ-slides (for live-cell experiments) (ibidi) and cultivated overnight at 23°C in HL-5 medium supplemented with 200 µM sodium palmitate. Infections (MOI 5) were performed with early stationary phase cultures of *L. pneumophila* JR32, Δ*icmT* or Δ*legG1* mutant strains harboring pNP99 (mCerulean), diluted in HL-5, and synchronized by centrifugation (450 *g*, 10 min, RT) (Rothmeier et al., 2013). Subsequently, infected cells were washed three times with HL-5 and incubated at 25°C for the time indicated. Finally, infected amoebae were recovered from the 6-well plates, fixed with 4% PFA for 30 min at RT, stained with LipidTOX Deep Red (1:1000 in SorC; Thermo-Fisher) for 30 min in the dark, transferred to 8-well µ-slides and embedded under a layer of PBS/0.5% agarose before imaging. For live-cell experiments, amoebae were stained prior to infection with LipidTOX Deep Red (1:200 in LoFlo medium) for 30 min in the dark and were directly imaged in LoFlo (ForMedium) in the 8-well µ-slides after washing.

Confocal microscopy of bacteria grown in liquid culture, fixed or live infected cells and LCV-LD *in vitro* was performed as described (Finsel et al., 2013; Rothmeier et al., 2013; Steiner et al., 2017; Weber et al., 2018) using a Leica TCS SP8 X CLSM with the following setup: white light laser, 442 nm diode and HyD hybrid detectors used for each channel. Pictures were taken using a HC PL APO CS2 63×/1.4 oil objective with Leica Type F immersion oil and analyzed with Leica LAS X software. Settings for fluorescence imaging were as following: DAPI (excitation 350 nm, emission 470 nm), mCerulean (excitation 442 nm, emission 469 nm), eGFP (excitation 488 nm, emission 516 nm), mCherry (excitation 568 nm, emission 610 nm), LipidTOX Deep Red (excitation 637 nm, emission, 655 nm). Images and movies were captured with a pinhole of 1.19 Airy Units (AU) and with a pixel/voxel size at or close to the instrument’s Nyquist criterion of approximately 43×43×130 nm (xyz). Scanning speed of 400 Hz, bi-directional scan and line accumulation equal 2 were used to capture still images (**Fig. 1**, **Fig. 3-5**, **Fig. S2A**, **Fig. S4-S5**). Resonant scanner (scanning speed: 8000 Hz) and line average equal 4 were used to capture movies (**Fig. 2**, **Fig. S2B**, **Supplementary Movies S1-S4**).

For image processing, all images and movies were deconvolved with Huygens professional version 19.10 (Scientific Volume Imaging, The Netherlands, http://svi.nl) using the CMLE algorithm with 40 iterations and 0.05 quality threshold. Signal to noise ratios were estimated from the photons counted for a given image. Single images, Z-stacks and movies were finalized and exported with Imaris 9.5.0 software (Bitplane, Switzerland) and analyzed using ImageJ software (https://imagej.nih.gov/ij/).

### Electron microscopy

Infection assays for electron microscopy were performed with *D. discoideum* Ax3 grown directly on grids as previously described (Medeiros et al., 2018). Briefly, the amoebae (5×10^5^ per well) were seeded onto EM gold finder grids (Au NH2 R2/2, Quantifoil) and incubated for 1 h, to allow the cells to attach to the grids. Cells were infected at an MOI of 100 and were vitrified at different time points (30 min and 3 h p.i.).

The vitrification of infected host cells was done by plunge-freezing as described previously (Weiss et al., 2017; Medeiros et al., 2018). In short, gold finder grids (Au NH2 R2/2, Quantifoil) containing infected amoebae were vitrified by back-blotting (Teflon, 2×3-7 sec). All grids were plunge-frozen in liquid ethane-propane (37%/63% v/v) using a Vitrobot (Thermo-Fisher) and stored in liquid nitrogen until further use.

Cryo-focused ion beam (cryoFIB) milling was used to prepare samples of plunge-frozen infected amoebae for imaging by cryo-electron tomography (cryoET) (Marko et al., 2007). Frozen grids with infected cells were processed as previously described (Medeiros et al., 2018) using a Helios NanoLab600i dual beam FIB/SEM instrument (Thermo-Fisher). Briefly, lamellae with ∼2 µm thickness were generated first (at 30 kV and ∼400 pA). The thickness of the lamellae (final ∼200 nm) was then gradually reduced using decreasing ion beam currents (final ∼25 pA). Lastly, lamellae were examined at low voltage by cryo-electron microscopy (cryoEM) imaging (3 kV, ∼0.17 nA) to visualize intracellular bacteria. CryoFIB-processed grids were unloaded and stored in liquid nitrogen until further use.

Frozen grids were examined by cryoEM and cryoET (Weiss et al., 2017). Data were collected on a Titan Krios TEM (Thermo-Fisher) equipped with a Quantum LS imaging filter and K2 Summit (Gatan). The microscope was operated at 300 kV, and the imaging filter was set to a slit width of 20 eV. The pixel size at the specimen level ranged from 3.45-5.42 Å. Tilt series of the lamellae were recorded from -60° to +60° with 2° increments and -8 µm defocus. The total dose of a tilt series was 75 e^-^/Å^2^ (intracellular *L. pneumophila*). Tilt series and 2D projection images were acquired automatically using SerialEM (Mastronarde, 2005). Three-dimensional reconstructions and segmentations were generated using the IMOD program suite (Mastronarde, 2005).

### Comparative proteomics of purified LDs

StrataClean beads for protein precipitation were incubated in 37% HCl for 6 h at 100°C, centrifuged (3’500 *g*, 5 min, RT) and washed twice to remove residual HCl. To extract LDs-bound proteins, LDs purified by density gradient centrifugation were lysed by freeze/thawing and by sonication (50% power, 3 s pulse for 2 min, 4°C). Extracted LDs protein sample solutions were then incubated with StrataClean beads in an over-head shaker for 2 h at 4°C. Following incubation, the beads were centrifuged (10’000 *g*, 45 min, 4°C) and the resulting pellet was washed once in distilled water (16’229 *g*, 10 min, 4°C). After supernatant removal, the bead pellet was dried in a vacuum centrifuge for 30 min and stored at 4°C.

StrataClean-bound LDs proteins were resolved by 1D-SDS-PAGE, the gel lanes were excised in ten equidistant pieces and subjected to trypsin digestion (Bonn et al., 2014). For the subsequent LC-MS/MS measurements, the digests were separated by reversed phase column chromatography using an EASY nLC 1200 (Thermo-Fisher) with self-packed columns (OD 360 μm, ID 100 μm, length 20 cm) filled with 3 µm diameter C18 particles (Dr. Maisch, Ammerbuch-Entringen, Germany) in a one-column setup. Following loading/ desalting in 0.1% acetic acid in water, the peptides were separated by applying a binary non-linear gradient from 5-53% acetonitrile in 0.1% acetic acid over 82 min. The LC was coupled online to a LTQ Orbitrap Elite mass spectrometer (Thermo-Fisher) with a spray voltage of 2.5 kV. After a survey scan in the Orbitrap (r = 60’000), MS/MS data were recorded for the twenty most intensive precursor ions in the linear ion trap. Singly charged ions were not considered for MS/MS analysis. The lock mass option was enabled throughout all analyzes.

After mass spectrometric measurement, database search against a database of *D. discoideum* downloaded from dictyBase (http://dictybase.org/) on 29/04/2022 (12,321 entries) as well as label-free quantification – LFQ (Cox et al., 2014) and iBAQ (Schwanhausser et al., 2011) – was performed using MaxQuant (version 1.6.7.0) (Cox and Mann, 2008). Common laboratory contaminants and reversed sequences were included by MaxQuant. Search parameters were set as follows: trypsin/P specific digestion with up to two missed cleavages, methionine oxidation and N-terminal acetylation as variable modification, match between runs with default parameters enabled. The FDRs (false discovery rates) of protein and PSM (peptide spectrum match) levels were set to 0.01. Two identified unique peptides were required for protein identification. LFQ was performed using the following settings: LFQ minimum ratio count 2 considering only unique for quantification.

Results were filtered for proteins quantified in at least two out of three biological replicates before statistical analysis. Here, both strains were compared by a student’s t-test applying a threshold p-value of 0.01, which was based on all possible permutations. Proteins were considered to be differentially abundant if the log2LFQ-fold change was greater than |0.8|. The dataset was also filtered for so-called “on-off” proteins. These proteins are interesting candidates as their changes in abundance might be so drastic that their abundance is below the limit of detection in the “off” condition. To robustly filter for these proteins, “on” proteins were defined as being quantified in all three biological replicates of the Δ*sey1* mutant setup and in none of the replicates of the Ax3 wild-type setup, whereas “off“ proteins were quantified in none of the three biological replicates of the mutant setup but in all replicates of the wild-type setup. In addition, the dataset was also filtered for the 50 most abundant proteins in each sample (based on the corresponding iBAQ value). After filtering, we ended with a list of 192 proteins that were either differentially abundant on LDs from *D. discoideum* Ax3 (39 proteins) or Δ*sey1* (76 proteins), or that belonged to the “on” (7 proteins) / “off” (22 proteins) categories or to the 50 most abundant proteins (48 additional proteins) (**Table S1**).

We then functionally mapped these proteins manually according to dictyBase classification, resulting in 135 functionally assigned and 57 unassigned proteins, and generated Voronoi treemaps (Paver v2.1, Decodon) to visualize functional clusters and abundance differences between LDs from *D. discoideum* Ax3 and Δ*sey1*. Individual iBAQ values of proteins on *D. discoideum* Ax3 LDs were represented by the area of the mosaic tiles (large tiles: more abundant, small tiles: less abundant). Comparison of protein abundance on Ax3 LDs with abundance of proteins on Δ*sey1* LDs is represented by the color gradient of the mosaic tiles (orange: more abundant on Ax3 LDs, grey: equally abundant on Ax3 and Δ*sey1* LDs, blue: more abundant on Δ*sey1* LDs).

### Isotopologue profiling

To label *L. pneumophila* intracellularly growing in *D. discoideum* with ^13^C-substrates, we used previously published protocols with minor modifications (Heuner and Eisenreich, 2013; Heuner et al., 2019). In brief, *D. discoideum* Ax3 or Δ*sey1* mutant producing calnexin-GFP (CnxA-GFP) were cultivated in eight T75 cell culture flasks per bacterial strain. After the cells reached confluency (∼2×10^7^ cells per flask), the amoebae were infected with *L. pneumophila* JR32 or Δ*fadL* (MOI 50), by adding bacteria grown in AYE to an OD_600_ of 5 at appropriate dilutions. The flasks were then centrifuged to synchronize infection (500 *g*, 10 min) and incubated for 1 h at 25°C. To remove extracellular bacteria, cells were washed once with 10 ml SorC buffer (room temperature), overlaid newly with 10 ml SorC buffer and further incubated at 25°C. At 5 h p.i., 200 µM [U-^13^C_16_]palmitate was added to the flasks, and the cells were further incubated for 10 h. At 15 h p.i., the amoebae were detached using a cell scraper, transferred into 50 ml Falcon tubes, frozen at -80°C for 1 h and again thawed to RT. The suspension was then centrifuged at 200 *g* for 10 min at 4°C, and the supernatant was transferred to new 50 ml Falcon tubes. The pellet, representing eukaryotic cell debris (F1) was washed twice with 50 ml and once with 1 ml cold SorC buffer. The supernatant harvested after the first centrifugation step contained *L. pneumophila* bacteria (F2). This fraction was centrifuged at 3500 *g* for 15 min at 4°C, and the resulting pellet was washed twice with 50 ml and once with 1 ml cold SorC buffer. The supernatant of F2 was filtered through a 0.22 µm pore filter to remove bacteria, and then 100% trichloroacetic acid (TCA) was added to a final concentration of 10%. The supernatant was incubated on ice for 1 h and centrifuged at 4600 × *g* for 30 min at 4°C. The resulting pellet (F3) contained cytosolic proteins of *D. discoideum*. The pellets of F1, F2 and F3 were autoclaved at 120°C for 20 min, freeze-dried and stored at -20°C until analysis. To monitor cell lysis and the purity of F1 and F2, samples were analyzed by light microscopy.

For isotopologue profiling of amino acids and polyhydroxybutyrate (PHB), bacterial cells (approximately 10^9^) or 1 mg of the freeze-dried host protein fraction were hydrolysed in 0.5 ml of 6 M HCl for 24 h at 105°C, as described earlier (Eylert et al., 2010). The HCl was removed under a stream of nitrogen, and the remainder was dissolved in 200 µl acetic acid. The sample was purified on a cation exchange column of Dowex 50Wx8 (H^+^ form, 200-400 mesh, 5 × 10 mm), which was washed previously with 1 ml methanol and 1 ml ultrapure water. The column was eluted with 2 ml distilled water (eluate 1) and 1 ml 4 M ammonium hydroxide (eluate 2). An aliquot of the respective eluates was dried under a stream of nitrogen at 70°C. The dried remainder of eluate 2 was dissolved in 50 µl dry acetonitrile and 50 µl *N*-(tert-butyldimethylsilyl)-*N*-methyltrifluoroacetamide containing 1% tert-butyldimethylsilyl chloride (Sigma) and kept at 70°C for 30 min. The resulting mixture of tert-butyldimethylsilyl derivates (TBDMS) of amino acids was used for further GC/MS analysis. Due to acid degradation, the amino acids tryptophan and cysteine could not be detected with this method. Furthermore, the hydrolysis condition led to the conversion of glutamine and asparagine to glutamate and aspartate. PHB was hydrolysed to its monomeric component 3-hydroxybutyric acid (3-HB). For derivatization of 3-HB, the dried aliquot of eluate 1 was dissolved in 100 µl *N*-methyl-*N*-(trimethylsilyl)-trifluoroacetamide (Sigma) and incubated in a shaking incubator at 110 rpm (30 min, 40°C). The resulting trimethylsilyl derivative (TMS) of 3-HB derived from PHB was used for GC/MS analysis without further treatment.

GC/MS-analysis was performed with a QP2010 Plus gas chromatograph/mass spectrometer (Shimadzu) equipped with a fused silica capillary column (Equity TM-5; 30 m × 0.25 mm, 0.25 µm film thickness; SUPELCO) and a quadrupole detector working with electron impact ionization at 70 eV. A volume of 0.1–6 µl of the sample was injected in 1:5 split mode at an interface temperature of 260°C and a helium inlet pressure of 70 kPa. With a sampling rate of 0.5 s, selected ion monitoring was used. Data was collected using LabSolution software (Shimadzu). All samples were measured three times (technical replicates). ^13^C-Excess values and isotopologue compositions were calculated as described before (Eylert et al., 2008) including: (i) determination of the spectrum of unlabeled derivatized metabolites, (ii) determination of mass isotopologue distributions of labeled metabolites and (iii) correction of ^13^C-incorporation concerning the heavy isotopologue contributions due to the natural abundances in the derivatized metabolites. For analysis of amino acids, the column was kept at 150°C for 3 min and then developed with a temperature gradient of 7°C min^-1^ to a final temperature of 280°C that was held for 3 min. The amino acids alanine (6.7 min), aspartate (15.4 min), glutamate (16.8 min) were detected, and isotopologue calculations were performed with m/z [M-57]^+^ or m/z [M-85]^+^. For the detection of 3-HB derived from PHB, the column was heated at 70°C for 3 min and then developed with a first temperature gradient of 10°C min^-1^ to a final temperature of 150°C. This was followed by a second temperature gradient of 50°C min^-1^ to a final temperature of 280°C, which was held for 3 min. The TMS-derivative of 3-HB, was detected at a retention time of 9.1 min, and isotopologue calculations were performed with m/z [M-15]^+^.

## Statistical analysis

If not stated otherwise, data analysis and statistics were performed in GraphPad Prism (Version 5.01, GraphPad Software Inc.) using two-way ANOVA with Bonferroni post-test. Probability values of less than 0.05, 0.01, and 0.001 were used to show statistically significant differences and are represented with *, **, or ***, respectively. The value of “n” represents the number of analyzed cells, LCVs, LDs or video frames per condition. For comparative proteomics, LFQ values were used for statistical testing of differentially abundant proteins. Empirical Bayes moderated t-tests were applied, as implemented in the R/Bioconductor limma package.

## Data availability

The MS proteomics data discussed in this publication have been deposited to the ProteomeXchange Consortium via the PRIDE (Perez-Riverol et al., 2019) partner repository with the dataset identifier PXD038200 (Reviewer account details: Username; Password, shkNQvEh).

## Abbreviations

Atl: atlastin
cryoET: cryo-electron tomography
Icm/Dot: intracellular multiplication/defective organelle trafficking
LCV: *Legionella*-containing vacuole
LD: lipid droplet
GFP: green fluorescent protein
MCS: membrane contact site
OSBP: oxysterol binding protein
PI: phosphoinositide
p.i.: post infection
Plin: perilipin
T4SS: type IV secretion system
VAMP: vesicle-associated membrane protein
Vap: VAMP-associated protein

## Acknowledgements

We would like to thank Caroline Barisch and Thierry Soldati for providing the GFP-Plin expression construct and the LDs isolation protocol. We would also like to thank Sebastian Grund for technical support during sample preparation before MS analyses. The Center for Microscopy and Image Analysis of the University of Zürich is acknowledged for maintaining and providing equipment, and ScopeM is acknowledged for instrument access at the ETH Zürich. Work in the group of H.H. was supported by the Swiss National Science Foundation (SNF; 31003A_175557, 310030_207826). The lab of M.P. was supported by the NOMIS foundation.

## Figure Legends

**Figure S1. Palmitate promotes intracellular replication of *L. pneumophila* in *D. discoideum* and increases lipid droplet numbers.** (**A**) *D. discoideum* Ax3, untreated (LoFlo medium) or treated with increasing concentrations of sodium palmitate (100-800 µM, 3 h), were infected (MOI 10) with GFP-producing *L. pneumophila* JR32 or Δ*icmT* (pNT28). The GFP-fluorescence was measured with a microtiter plate reader at 1 h, 24 h and 48 h p.i. Data show the relative fluorescence increase between 1 h and 24 or 48 h p.i. (JR32: black/grey bar; Δ*icmT*: white/dotted bar). Data represent means ± SD of three independent experiments (\**p* < 0.05). (**B, C**) Quantification of (B) LD number or (C) LD area in unstimulated or sodium palmitate stimulated (200 µM, overnight) *D. discoideum* Ax3 and Δ*sey1* producing AmtA-GFP (pHK121) and mCherry-Plin (pHK102), fixed with PFA, and stained with LipidTOX Deep Red (n^LDs^ > 250). Data represent means ± SD of three independent experiments (\*\*\**p* < 0.001).

**Figure S2. Localization of Ran and RanBP1 to LDs and dynamics of LCV-LD interactions.** (**A**) Representative fluorescence micrographs of *D. discoideum* Ax3 producing RanA-mCherry (pLS221) and GFP-Plin (pHK101) (**top**) or RanBP1-GFP (pLS222) and mCherry-Plin (pHK102) (**bottom**). Scale bars: overview (2 µm), inset (1 µm). (**B**) Representative fluorescence micrographs of *D. discoideum* Ax3 or Δ*sey1* producing P4C-GFP (pWS034) and mCherry-Plin (pHK102), fed overnight with 200 µM sodium palmitate, stained with LipidTOX Deep Red and infected (MOI 5) with mCerulean-producing *L. pneumophila* JR32 (pNP99). Infected cells were recorded for 60 s each at the time indicated using a Leica TCS SP8. Single fluorescence channels of the 60 min p.i. recordings (Fig. 2A) are shown. Scale bars: 0.5 µm.

**Figure S3. Alignment of FadL homologues.** FadL homologues of *E. coli*, *L. pneumophila*, *V. cholerae*, and *P. aeruginosa* were aligned using the ClustalW algorithm available in the Clustal Omega web browser. The symbols underneath the alignment indicate the degree of conservation: identical residues (**\***), highly similar residues (:), similar residues (.). The red box denotes the strictly conserved NPA motif in the “hatch” domain (amino acids 73-75 in *L. pneumophila* FadL, Lpg1810).

**Figure S4. Expression of *L. pneumophila fadL* and growth of the Δ*fadL* mutant strain in medium.** (**A**) Representative fluorescent micrographs of *L. pneumophila* JR32, Δ*lqsA*, Δ*lqsR,* and Δ*lvbR* harboring P*_fadL_*-*gfp* (pPS003), grown in AYE broth at 37°C, fixed with PFA and stained with DAPI. Bacteria were imaged at the times indicated using a Leica TCS SP8 microscope. Representative images of three independent experiments. Scale bar: 10 μm. (**B**) *L. pneumophila* JR32, Δ*lqsA*, Δ*lqsR,* or Δ*lvbR* mutant strains harboring P*_fadL_*-*gfp* (pPS003) or JR32 harboring P*_flaA_*-*gfp* (pCM09) were grown in AYE broth at 37°C. OD_600_ and GFP fluorescence (relative fluorescence units, RFUs) and OD_600_ (microplate reader) were measured over 48 h. Data represent means ± SD of three independent experiments (\*\*\**p* < 0.001). (**C**) The *pneumophila* parental strain JR32 (**circles**) and Δ*fadL* (**squares**) were grown at 37°C in AYE (**black**) or MDM (**violet**), and OD_600_ (microplate reader) was monitored for 48 h. Data represent means ± SD of three independent experiments.

**Figure S5. Intracellular *L. pneumophila* expresses *fadL* in *D. discoideum*.** (**A**) Confocal microscopy of *D. discoideum* infected (MOI 5) for the time indicated with *L. pneumophila* JR32 harboring P*_fadL_*-*gfp* (pPS003). The cells were fixed in 4% PFA, permeabilized with ice-cold methanol, and stained with 1 µg/ml DAPI. Images shown are representative of two independent experiments. Scale bars: 5 μm. (**B**) Quantification of GFP-positive *L. pneumophila* JR32 (pPS003) growing in *D. discoideum*. At the time p.i. indicated, host cells were lysed with 0.1% Triton X-100, fixed in 4% PFA, stained with 1 µg/ml DAPI and analyzed by flow cytometry and the software FlowJo. Plotted are means and individual data points of two independent experiments.

**Table S1. Comparative proteomics analysis of LDs from D. discoideum Ax3 or Δsey1 mutant amoeba.** Dark blue: exclusively present on Δsey1 LDs, light blue: more abundant on Δsey1 LDs, grey: equally abundant on Ax3 and Δsey1 LDs, light orange: more abundant on Ax3 LDs, dark orange: exclusively present on Ax3 LDs.

**Table S2. Cells, bacterial strains, and plasmids used in this study.**

**Table S3. Oligonucleotides used in this study.**

**Supplementary Movie S1**

Representative movie of *D. discoideum* Ax3 producing P4C-GFP (pWS034) and mCherry-Plin (pHK102), fed overnight with 200 µM sodium palmitate, stained with LipidTOX Deep Red and infected (MOI 5) with mCerulean-producing *L. pneumophila* JR32 (pNP99). Infected cells were recorded for 60 s each at the times indicated using a Leica TCS SP8.

**Supplementary Movie S2**

Representative movie of *D. discoideum* Ax3 producing P4C-GFP (pWS034) and mCherry-Plin (pHK102), fed overnight with 200 µM sodium palmitate, stained with LipidTOX Deep Red and infected (MOI 5) with mCerulean-producing *L. pneumophila* Δ*legG1* (pNP99). Infected cells were recorded for 60 s each at the times indicated using a Leica TCS SP8.

**Supplementary Movie S3**

Representative movie of *D. discoideum* Δ*sey1* producing P4C-GFP (pWS034) and mCherry-Plin (pHK102), fed overnight with 200 µM sodium palmitate, stained with LipidTOX Deep Red and infected (MOI 5) with mCerulean-producing *L. pneumophila* JR32 (pNP99). Infected cells were recorded for 60 s each at the times indicated using a Leica TCS SP8.

**Supplementary Movie S4**

Representative movie of *D. discoideum* Δ*sey1* producing P4C-GFP (pWS034) and mCherry-Plin (pHK102), fed overnight with 200 µM sodium palmitate, stained with LipidTOX Deep Red and infected (MOI 5) with mCerulean-producing *L. pneumophila* Δ*legG1* (pNP99). Infected cells were recorded for 60 s each at the times indicated using a Leica TCS SP8.

## Notes

### Competing Interest Statement

The authors have declared no competing interest.

